# Blindness reshapes mental time travel: From perceptual scenes to conceptual scaffolds

**DOI:** 10.64898/2026.03.27.714680

**Authors:** Nadja Abdel Kafi, Marie Malinowski, Pitshaporn Leelaarporn, Julia Taube, Christine Kindler, Marion Crump, Anja Essmann, Sven Lange, Sarah Dumitrescu, Nicola Mattar, Ella Gutenberg, Sascha Brunheim, Tony Stöcker, Katharina Wall, Bettina Wabbels, Annika Spottke, Cornelia McCormick

## Abstract

How can humans remember and imagine without vision? Mental time travel, the ability to re-experience past events and envision future ones, is widely assumed to rely on visual imagery and the construction of mental scenes. Blindness provides a critical test of this assumption. Across behavioral interviews, language analyses, and multimodal neuroimaging in congenitally blind, late-blind, and sighted individuals, we show that blind individuals, even those blind from birth, mentally time travel as vividly as sighted people, but construct their inner worlds differently. Sighted participants relied on perceptual detail and activated classic scene-processing regions, whereas blind participants emphasized thoughts and emotions and recruited reorganized occipital cortex. Connectivity analyses revealed strengthened coupling between occipital and medial temporal regions, indicating adaptive reconfiguration of the episodic system. The brain does not require images to imagine: it flexibly builds internal experiences using the representational resources available.

**Summary Paragraph:** How humans reconstruct events that are no longer available to the senses is a fundamental but unresolved question. Remembering the past and imagining the future, known as mental time travel, is a defining feature of human cognition, shaping identity and guiding decisions¹,². Prevailing theories assume that such constructions depend on visual imagery, with the mind’s eye reconstructing events on a visuospatial stage³,⁴. Blindness provides a critical test of this assumption. Across extensive behavioral interviews and multimodal neuroimaging, we find that mental time travel remains phenomenologically intact in blindness, but its scaffolding changes fundamentally. Sighted individuals rely on perceptual detail and activate regions specialized for visual scenes, whereas blind individuals, whether blind from birth or later in life, emphasize thoughts and emotions and reorganize occipital cortex for conceptual strategies⁵,⁶. These contrasting strategies map onto distinct neural signatures, revealing a dissociation between perceptual and conceptual routes to episodic simulation. Together, these findings reveal that the brain’s capacity to construct internal experience rests on conceptual scaffolding, not perceptual re-creation⁷.

## Introduction

How do humans reconstruct worlds that are no longer available to the senses? Whether we are reliving a childhood moment or imagining a future possibility, we effortlessly summon internal experiences that feel rich, coherent, and personally meaningful. This capacity, often described as mental time travel, depends on a distributed brain network centered on the hippocampus¹,². This system constructs richly detailed mental scenes that integrate sensory, spatiotemporal, and emotional elements, enabling episodic memory and imagination^4,8^. In sighted individuals, scene construction typically relies on visuospatial scaffolds grounded in visual imagery^9,10^. In fact, visualization abilities predict the vividness and detailedness of autobiographical memory (AM) and future thinking^3,11^. Furthermore, impairments in mental visualization, such as in aphantasia, are associated with reduced episodic detail and diminished reliving^12–14^. Together, these findings have led to the wide-spread assumption that visual imagery is indispensable for vivid, detail-rich mental time travel.

Blindness offers a natural and powerful test of this assumption. If visual experience is essential for mental time travel, individuals who are blind, particularly those blind from birth, should exhibit profound deficits in episodic simulation. Yet empirical evidence is strikingly scarce. Behavioral studies in blind individuals suggest that autobiographical memory remains largely intact, even though these individuals construct memories using non-visual scaffolds such as auditory, and tactile details^15–17^. Some findings even indicate superior auditory memory performance in blindness^6,18^, hinting at the possibility that the episodic system reorganizes when visual input is absent. However, no prior work has examined mental time travel combining behavioral and neuroimaging approaches. In addition, the few studies available are heterogeneous, and rarely differentiate congenital from late blindness, a critical distinction given the developmental role of early visual input in shaping cortical architecture^19,20^.

These issues are central to the constructive framework of episodic memory developed by Eleanor Maguire and colleagues, whose work established scene construction as a core computational process of mental time travel^4,8,21,22^. Building on this theoretical legacy, we asked whether such constructive processes require visual experience, or whether the episodic system can operate through alternative, conceptual scaffolds.

Here, we address this gap with a multimodal study of congenitally blind, late-blind, and sighted individuals. We hypothesized that blindness would mimic aphantasia, leading to reduced visual imagery and diminished mental time travel. To test this, we combined structured interviews of autobiographical memory, scene construction, and episodic future thinking with functional MRI, connectivity analyses, and diffusion imaging. This approach allowed us to examine whether visual experience is essential for constructing mental events and to identify the behavioural and neural strategies that support episodic simulation when vision is absent.

## Results

### Participants and analysis approach

We studied 81 adults: 22 congenitally blind (CB), 22 late-blind (LB), and 37 sighted controls (CTL). All blind participants were completely blind as confirmed by ophthalmological examination; a few retained minimal light sensitivity but could not perceive shapes or distinguish sky from ground. In the LB group, blindness onset occurred at a mean age of 31 years (SD = 20), and the mean duration of blindness was 29 years (SD = 20). Groups were matched on sex and education but differed in age: LB participants were older than CB (p = 0.04), whereas CTL did not differ from either blind group. Supplementary materials provide full details on causes of blindness, demographic profiles, recruitment procedures, ethical approval, and cognitive measures (MoCA, BDI). All group analyses were adjusted for age using ANCOVA or MANCOVA models.

### Imagery vividness

Self-report measures revealed clear group differences in visual imagery vividness (Vividness of Visual Imagery Questionnaire, VVIQ^23^; Fig. 1a; ANCOVA, p < 0.001; full statistics in Supplementary Methods). Sighted controls showed the highest vividness, followed by LB and then CB. Importantly, CB participants did not classify themselves as aphantasic: their scores were significantly above the standard cut-off of 32 (one-sample t-test, p = 0.007). On the Plymouth Sensory Imagery Questionnaire (PSIQ)^24^, overall imagery did not differ across groups (p = 0.68). However, modality-specific analyses showed that CB participants reported markedly reduced visual imagery compared with LB and CTL (p’s < 0.001), whereas non-visual modalities were comparable across groups. Together, these findings indicate that non-visual imagery is preserved across all groups, whereas visual imagery is specifically and markedly reduced in CB.

**Figure 1.**
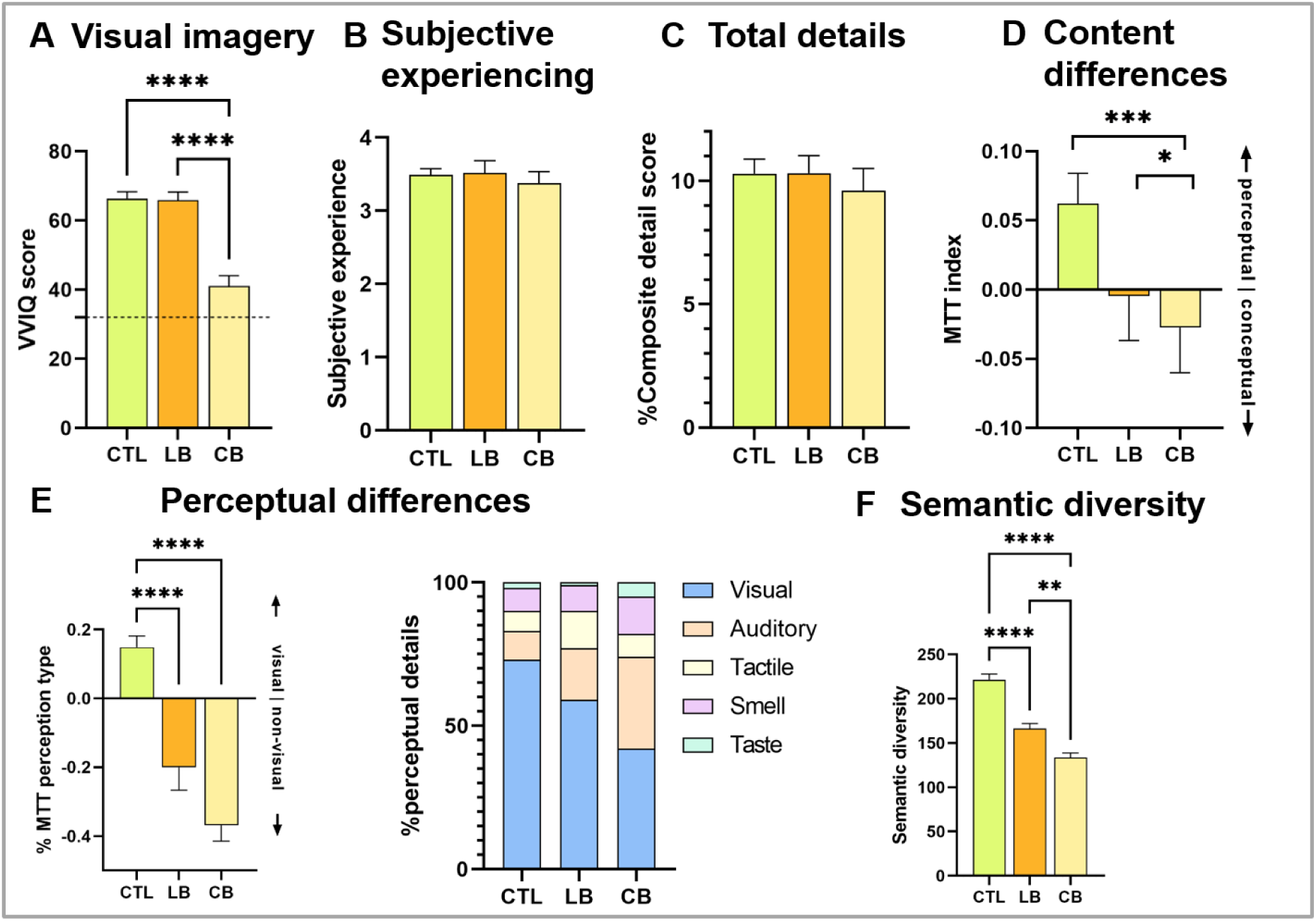
Mental time travel profiles differ by visual experience. (a) Vividness of Visual Imagery Questionnaire (VVIQ) scores were highest in sighted controls (CTL), intermediate in late-blind (LB), and lowest in congenitally blind participants (CB); the broken line marks the aphantasia threshold (score = 32). CB participants scored significantly above this threshold (p < 0.01). (b) Subjective ratings of mental time travel did not differ between groups. (c) Total detail density across autobiographical memory, scene construction, and episodic future thinking interviews did not differ between groups. (d) Composite mental time travel (MTT) scores revealed perceptual dominance in CTL and conceptual dominance in CB, with LB showing an intermediate profile. (e) Sensory composition of perceptual details differed systematically: CTL relied primarily on visual details, CB on non-visual modalities (auditory, tactile), and LB showed mixed profiles. The stacked bar graph illustrates visual dominance in CTL and non-visual reliance in CB. (f) Lexical diversity differed significantly across groups: CTL produced the most diverse and idiosyncratic descriptions, LB intermediate, and CB the least, indicating more structured and repetitive conceptual scaffolds in congenital blindness. Bars show means ± SEM.

### Mental time travel shifts from perceptual to conceptual

If visual imagery is diminished in congenital blindness, does mental time travel suffer? Subjective ratings and the overall detailedness of imagined or recalled events were comparable across groups, yet their content diverged markedly (Fig. 1b,c). CB participants provided significantly more conceptual details, such as thoughts and emotions, whereas CTL relied predominantly on perceptual details. LB showed an intermediate pattern. A composite mental time travel (MTT) score contrasting perceptual and conceptual detail densities confirmed this gradient: CTL scored positively, indicating perceptual dominance, whereas CB scored negatively, reflecting conceptualization (p < .001; Fig. 1d). Because autobiographical memory^25^, scene construction⁸, and episodic future thinking interviews showed highly similar patterns (Supplementary Methods; Figures S1–S3), we concatenated these measures to compute composite scores.

### Blindness reshapes sensory composition of mental time travel

When perceptual details were broken down by sensory modality, CTL narratives were dominated by visual descriptions, whereas CB participants relied primarily on non-visual modalities, especially auditory, smell and tactile details. Notably, in the autobiographical memory interview, CB participants referred to “visual” details more often than auditory ones (paired t-test: t(20) = 2.584, p = .018), indicating the use of conceptual rather than perceptual visual terms despite lifelong blindness (Fig. 1e; Supplementary Methods; Figures S1–S3). The same directional pattern was present in scene construction and episodic future thinking and was preserved in the composite scores, although it did not reach significance outside the AM interview. LB participants showed mixed sensory profiles, consistent with partial retention of visual imagery.

### Semantic diversity is reduced in blindness

Congenitally blind participants relied more on conceptual than perceptual details. But how varied are these conceptual constructions? During qualitative coding, we noticed that descriptions, especially in scene construction, often followed highly similar linguistic templates across blind participants, with different individuals using nearly the same words to depict the same scenarios. For example, when describing the scene of lying at the edge of a swimming pool in a luxury hotel, congenitally blind participants often produced nearly identical formulations, typically a warm, sunny setting with a sunlounger, a soft towel, and a sun umbrella close to the pool, whereas sighted controls described a far broader range of scenarios (e.g., being in the water, chatting at a pool bar, or approaching the scene from a distance).To formally quantify this impression, we computed a semantic diversity measure that captures the range of unique words used to describe autobiographical and imagined events (Fig. 1f; Supplementary Methods). Sighted participants showed the greatest lexical diversity, late-blind individuals intermediate, and congenitally blind individuals the lowest (p < 0.001, η² = 0.58).

Together, these findings suggest that congenital blindness is associated with more structured and repetitive conceptual scaffolds during mental time travel, whereas sighted individuals draw on a broader and more varied semantic repertoire.

### Neural architecture of mental time travel in blindness

These behavioral results show that mental time travel remains intact in blindness but supported by different representational strategies. This motivates the central question for the neural analyses: how does the brain construct mental events without visual experience?

Participants successfully performed both autobiographical memory (AM) and scene construction (SC) tasks during fMRI scanning. Both paradigms used auditory cues, realistic event prompts for AM (e.g., “a party”) and unrealistic scenarios for SC (e.g., “being on the moon”), with math trials serving as controls. After each trial, participants rated vividness or difficulty, and performance did not differ between groups (all p’s > 0.2). Across tasks, ∼80% of trials were rated as vivid or easy. Full task details and separate AM/SC results are provided in the Supplementary Methods.

Task-based fMRI revealed a common episodic network across all groups, including bilateral hippocampus, parahippocampal cortex, posterior cingulate cortex (PCC), lateral parietal cortex, and ventromedial prefrontal cortex (vmPFC; Fig. 2a). Group differences emerged in regions associated with perceptual versus conceptual strategies: sighted participants showed stronger activation in scene-selective areas, including a parahippocampal place-related region (PPA-related)^26^ and fusiform gyrus, whereas congenitally blind participants exhibited greater engagement of PCC and lateral occipital cortex (OCC) bilaterally (ANCOVA with age as covariate; Fig. 2b,c). Because AM and SC produced highly similar activation patterns (Supplementary Figures S4–S5), we combined them for the main analysis.

**Figure 2.**
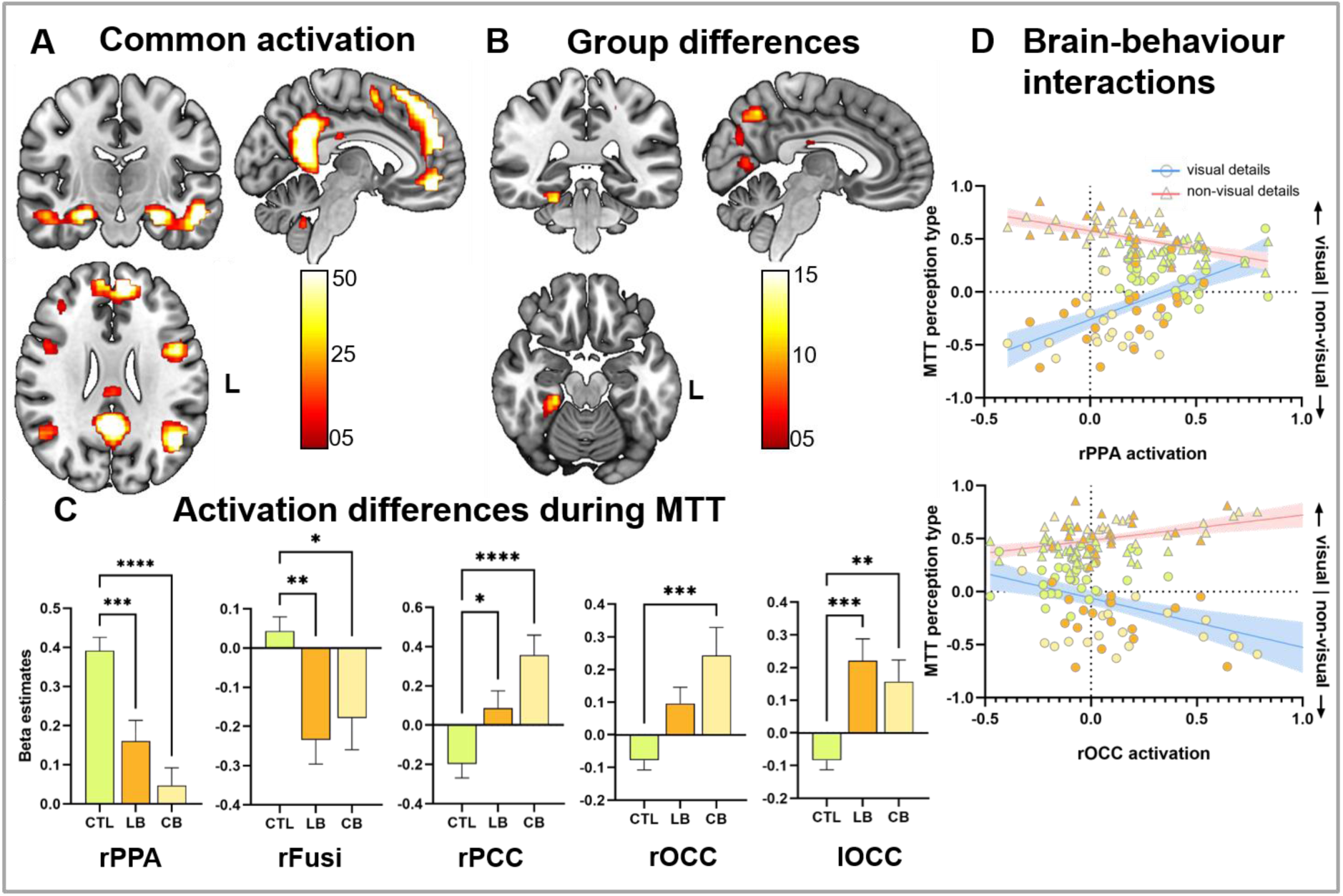
Neural architecture and group-specific adaptations in mental time travel. (a) Task-based fMRI during autobiographical memory and scene construction revealed a common episodic network across all groups, including bilateral hippocampus, parahippocampal cortex, posterior cingulate cortex (PCC), lateral parietal cortex, and ventromedial prefrontal cortex (vmPFC). (b) Group differences emerged in regions supporting mental time travel: sighted participants showed stronger activation in the parahippocampal place-related area (PPA-related) and fusiform gyrus, whereas congenitally blind participants exhibited greater engagement of PCC and lateral occipital cortex (LOC). (c) Beta estimates from regions showing significant group differences in (b), illustrating effect magnitude and direction. (d) Brain–behavior correlations revealed a double dissociation: PPA-related activation tracked visual-perceptual detail density and inversely with non-visual detail, whereas occipital activation showed the opposite pattern (see Fig. 1b), linking neural activity to content-specific strategies. Bars show means ± SEM; significance markers indicate ANCOVA results (****p < 0.0001; ***p < 0.001; **p < 0.01; *p < 0.05).

### Distinct neural signatures for perceptual and conceptual strategies

Across all groups, neural activity mirrored the behavioral shift from visual-perceptual to non-visual strategies, revealing a clear double dissociation (Fig. 2d; Supplementary Methods). A PPA-related region tracked visual-perceptual detail density (r = 0.42, p < 0.001) and inversely tracked non-visual detail (r = –0.38, p < 0.001), whereas right occipital cortex showed the opposite pattern, correlating positively with non-visual detail density and negatively with visual detail (all p’s < 0.05). Similar modality-specific patterns were observed in other regions associated with visual imagery (Fig. S6), including right fusiform cortex, bilateral hippocampus, and vmPFC. Right PCC and left OCC showed patterns comparable to right OCC, whereas a control region (dorsolateral prefrontal cortex) showed no relationship with either visual or non-visual detail (all p’s > 0.31). Together, these reciprocal associations in PPA-related and occipital regions indicate that perceptual and conceptual strategies rely on distinct neural substrates, an important hallmark of functional specialization.

### Rewiring the episodic system in blindness

Blindness alters interactions within the episodic system, strengthening functional coupling while reducing structural integrity. Because the PPA-related region showed the strongest group differences in activation, we examined its connectivity profile. Both resting-state and task-based analyses revealed increased functional coupling between the PPA-related region, occipital cortex, and fusiform gyrus in congenitally blind participants (Fig. 3a,b; Supplementary Methods; Fig. S7). In contrast, diffusion tensor imaging showed reduced fractional anisotropy (FA) in occipital white matter compared to sighted controls, indicating widespread structural reorganization (Fig. 3c; Supplementary Methods; Tables S3–S5). Within the congenitally blind group, higher FA values correlated positively with mental time travel scores (p < 0.05, TFCE-corrected), suggesting that preserved occipital integrity supports more perceptual-like episodic simulation.

**Figure 3.**
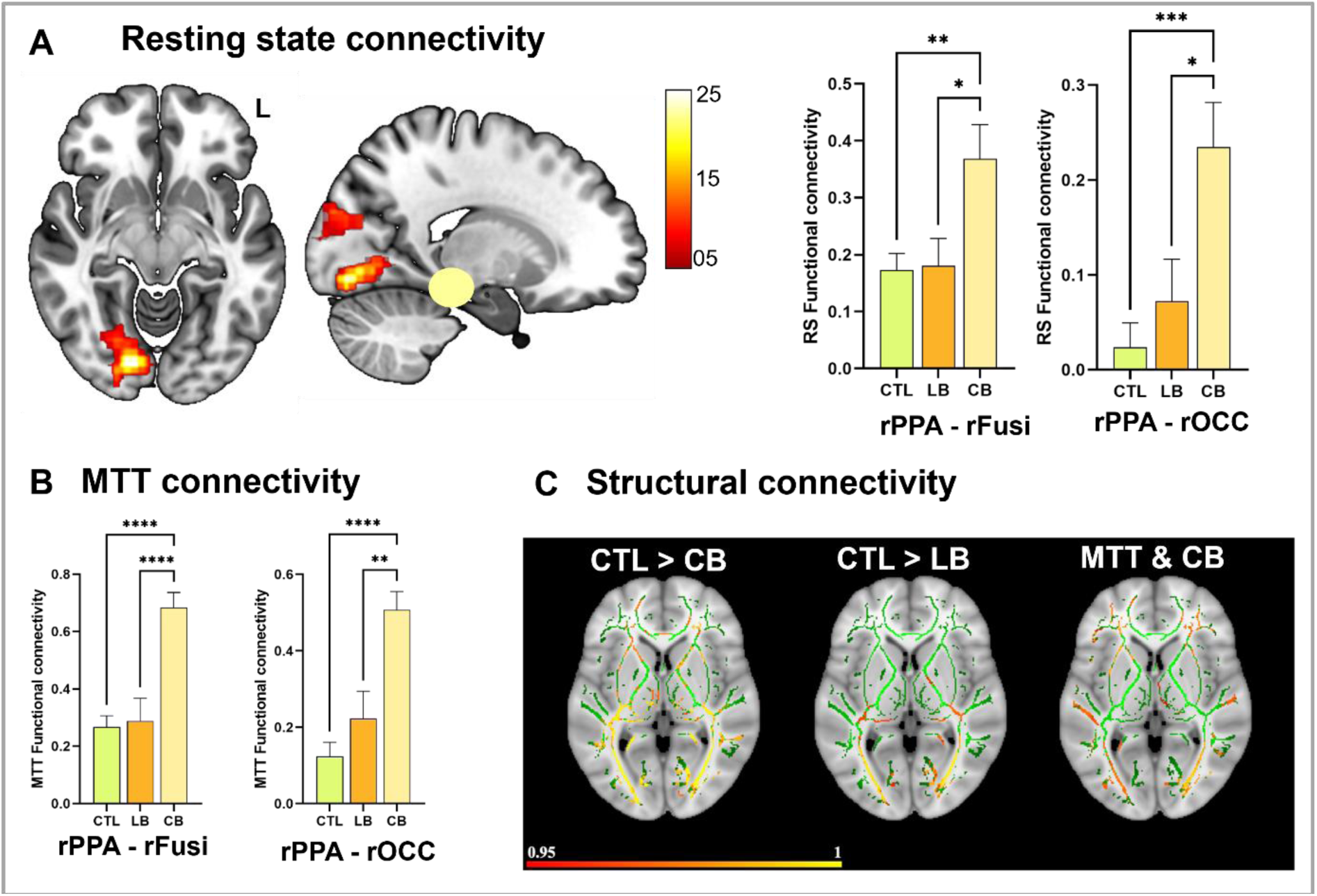
Functional connectivity and white matter integrity and adaptations in blindness. (a) Resting-state and (b) task-based connectivity analyses revealed increased coupling of the parahippocampal place-related area (PPA-related, yellow sphere) with occipital and fusiform cortex in congenitally blind participants compared to sighted controls. (c) Tract-based spatial statistics (TBSS) showed reduced fractional anisotropy (FA) in occipital white matter tracts in congenitally blind individuals relative to sighted and late-blind participants (CTL > CB; CTL > LB). Within the congenitally blind group, higher FA values were positively associated with mental time travel performance, indicating that preserved occipital integrity supports more perceptual-like episodic simulation. Statistical maps (red–yellow) represent TFCE-corrected results (p < 0.05) overlaid on the mean FA skeleton (green) in MNI152 space. Connectivity maps and bar graphs show means ± SEM; significance markers indicate ANCOVA results (****p < 0.0001; ***p < 0.001; **p < 0.01; *p < 0.05).

### Occipital grey-matter alterations with preserved hippocampal volume

Voxel-based morphometry and volumetric analyses revealed significant group differences in grey-matter volume, with CB and LB showing reduced grey matter particularly in occipital regions compared to CTL (Supplementary Methods; Fig. S8). In contrast, hippocampal volume did not differ between groups (p = 0.16). These results indicate that structural adaptations in blindness are primarily occipital and connectivity-based, rather than involving core episodic-memory regions such as the hippocampus.

### An integrated axis of mental time travel strategies

To test whether behavioral, functional, and structural measures converge on a shared underlying organization, we entered all variables into a single principal component analysis, without including group membership. This unsupervised approach revealed a dominant perceptual–conceptual axis that accounted for 31% of the variance (p < 0.0001; Fig. 4; Supplementary Methods). Loadings on the perceptual side included visual imagery vividness, perceptual detail density, stronger activation in the PPA-related region, and higher occipital white-matter integrity. Loadings on the conceptual side reflected reliance on non-visual detail, greater occipital activation, and stronger PPA–occipital coupling. Importantly, variables that did not show prior group differences, such as hippocampal volume and internal MTT density, also aligned meaningfully along this axis, underscoring its robustness. Although the PCA was performed oblivious to group identity, the resulting component scores showed a striking separation across groups, revealing a clear gradient (CTL > LB > CB, all p < 0.001). Thus, the data-driven axis integrates all modalities into a unified perceptual–conceptual continuum, showing how behavioral strategies, functional recruitment, and structural architecture systematically coalesce in the presence or absence of visual experience.

**Figure 4.**
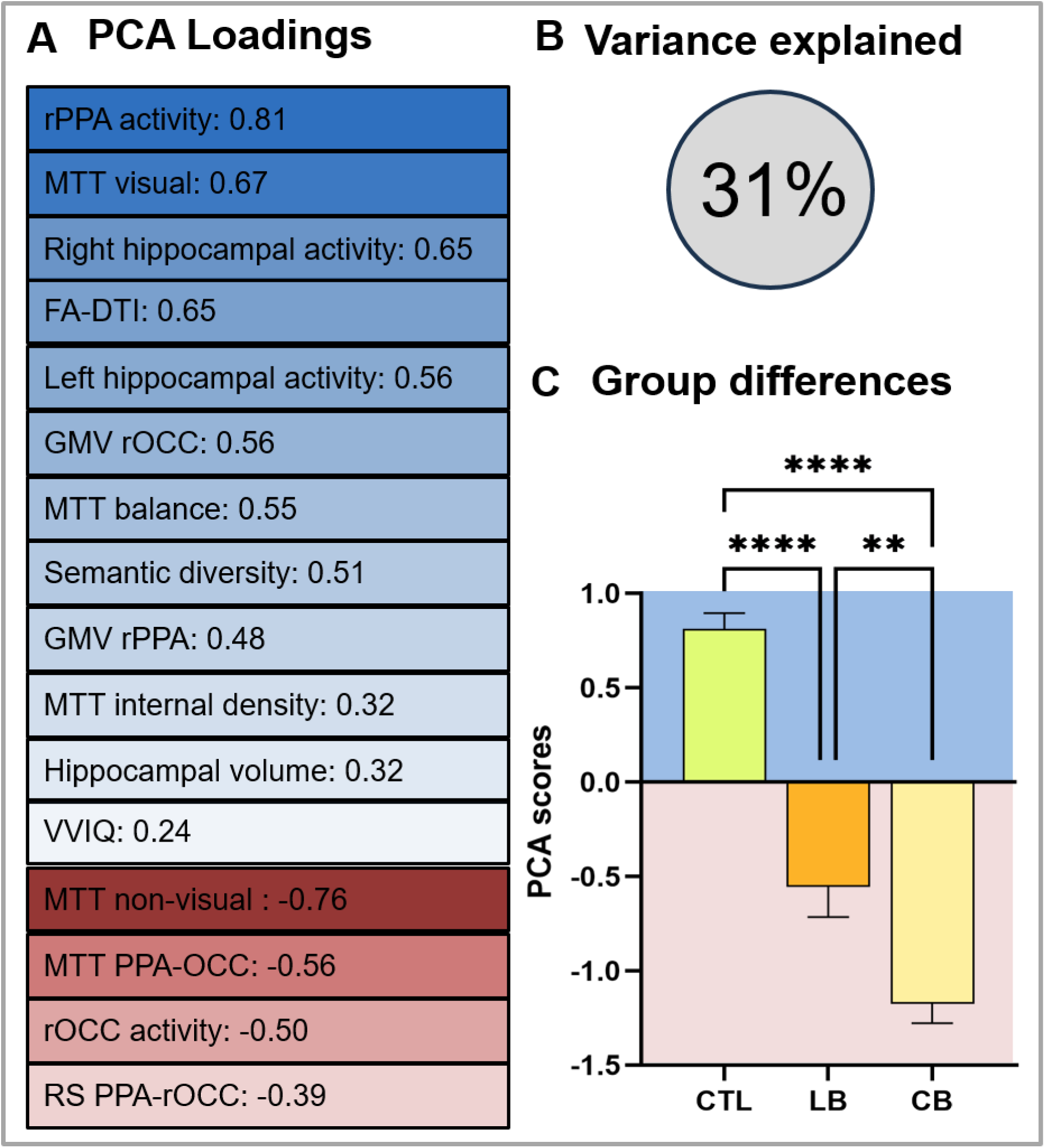
An integrated axis of mental time travel strategies. (a) A multimodal principal component analysis—including behavioral measures (imagery vividness, perceptual–conceptual detail balance) and neuroimaging indices (task-based activation, connectivity, fractional anisotropy)—revealed a dominant perceptual–conceptual axis summarizing mental time travel strategies. Positive loadings reflected visual imagery, perceptual detail, parahippocampal (PPA-related) activation, and higher occipital FA, whereas negative loadings reflected reliance on non-visual strategies, greater occipital activation, and stronger PPA–occipital connectivity. (b) This axis accounted for 39.3% of variance in the multimodal dataset (observed eigenvalue = 3.596; random-permutation mean = 1.720; p < 0.0001). (c) Although computed without group labels, component scores showed a clear gradient across groups (p < 0.0001, η² = 0.732), with sighted controls highest, late-blind intermediate, and congenitally blind lowest. Blue shading denotes perceptual mental time travel; red shading denotes conceptual mental time travel. Error bars indicate SEM.

## Discussion

### Perceptual–conceptual scaffolding in mental time travel

Our multimodal approach shows that mental time travel remains intact without visual experience, yet its underlying scaffolding and neural architecture shifts fundamentally. This finding challenges the long-standing assumption that visual imagery is required for episodic simulation^4,8^ and instead reveals a flexible, modality-independent system that adapts to available sensory resources.

In sighted individuals, autobiographical memory, scene construction, and episodic future thinking were consistently dominated by perceptual detail and accompanied by strong engagement of a PPA-related region^4,8^. Evidence from aphantasia further supports this link. Individuals who lack voluntary visual imagery recall fewer episodic details and show altered hippocampal–visual cortex connectivity^7,12–14^.

By contrast, congenitally blind participants, despite never having had visual experience, recalled and imagined events with comparable richness, relying on conceptual scaffolds such as thoughts and emotions rather than perceptual scenes. This perceptual-to-conceptual shift aligns with preliminary behavioral work showing that blind individuals retrieve autobiographical memories rich in emotional and semantic content^15–17^.

### Conceptual “visual” representations without visual experience

Congenitally blind participants frequently produced details that score as “visual,” particularly during autobiographical memory. One participant, for instance, described the ornate shape of a chandelier at his own wedding, information he could only have acquired second-hand. Such examples illustrate how conceptually learned event knowledge becomes woven into personal memory, blurring the line between external input and internal reconstruction. They also reflect a broader cultural reality. Because language and shared discourse are predominantly visual, blind individuals naturally adopt visual terminology to structure their event models.

Our scoring adhered to widely used autobiographical memory and scene-construction frameworks^8,25^, in which object-based descriptions (e.g., a small chair) are classified as visual for sighted populations. In congenitally blind individuals, however, these elements almost certainly reflect conceptual, not perceptual, visual representations. This distinction underscores a central insight of our study. Blind individuals do not merely substitute non-visual sensations; they construct events using conceptually “visual” elements, showing that visual terminology and structure can emerge without perceptual vision^27–30^. This phenomenon offers content-level evidence that event construction is anchored in conceptual scaffolding, rather than tied to a specific sensory modality.

### Occipital integration into the episodic system

The neural dissociation we observed suggests more than group-specific recruitment differences. It points to a reorganization of representational roles within the episodic system. While sighted individuals relied on perceptual scaffolding supported by a PPA-related region and fusiform cortex^31,32^, congenitally blind participants engaged occipital regions typically associated with vision and showed increased coupling between these areas and the PPA-related region. Rather than reflecting mere cross-modal compensation, this pattern may indicate that occipital cortex in congenital blindness^5,6,33^ becomes functionally integrated into the broader default-mode/autobiographical memory network^34–36^. Such integration could allow occipital cortex to support multimodal, non-visual representations that serve conceptual aspects of mental time travel. This interpretation aligns with our brain–behavior correlations, that activity in the PPA-related region tracked perceptual detail density, whereas occipital activation scaled with conceptual strategies. Under this view, the episodic system preserves its constructive capacity by reallocating representational functions, transforming occipital cortex into a conceptual processing node within the broader event-construction architecture.

### Hippocampal contributions to event construction

Across all groups, the hippocampus remained a central hub for mental time travel, with preserved volume and robust engagement^2,37^. In congenital blindness, this sustained involvement likely reflects a network-level reconfiguration in which repurposed occipital regions supply conceptual inputs that the hippocampus binds into coherent event models. In sighted individuals, hippocampal activity typically aligns with visual scene construction^38,39^, and our trend-level group effect is consistent with this bias. Crucially, brain–behavior correlations show that hippocampal responses track perceptual detail when available and shift away from conceptual content. These patterns suggest that the hippocampus is not inherently visual but preferentially anchors event construction to the richest representational format, perceptual in the sighted, conceptual in the blind.

### Developmental traces of visual imagery in late-blind individuals

Late-blind individuals offer a unique window into how visual experience shapes episodic construction. Behaviorally, they retained more perceptual detail than congenitally blind participants, likely reflecting residual visual representations from earlier life^16,17^, yet their profiles remained intermediate and vividness ratings did not differ reliably from the other groups. Neurally, they showed occipital reorganization and reduced structural integrity in visual pathways, but to a lesser degree than congenitally blind participants^5,6,33^. This pattern suggests two interacting influences: early visual experience leaves a lasting trace that can support perceptual richness, while subsequent plasticity gradually shifts the system toward non-visual strategies as visual input is lost. Together, these observations indicate that mental time travel is shaped not only by sensory modality but also by developmental history and the timing of visual loss, consistent with work on navigation and conceptual knowledge in blindness^27,28,40,41^.

### Beyond visual imagery: A conceptual route to episodic construction

Our findings refine models of episodic memory by showing that constructing coherent event models, rather than visual imagery per se, is the core computational process of mental time travel. While scene construction theory emphasizes visuospatial imagery in sighted individuals, and aphantasia research links reduced imagery to diminished episodic detail, blindness reveals a different route. Rich episodic simulation can emerge from non-visual scaffolds. This challenges the assumption that perceptual detail is necessary for vivid recall and supports a modality-independent architecture for event construction. The double dissociation between PPA-related and occipital regions further indicates that perceptual and conceptual strategies rely on distinct neural dimensions, illustrating how the episodic system adapts to sensory constraints. Finally, our multimodal PCA provides convergent evidence that these behavioral and neural signatures form a coherent perceptual–conceptual axis. Computed without group labels, the PCA revealed a dominant component on which visual imagery, perceptual detail, PPA-related activation, and occipital white-matter integrity loaded positively, whereas non-visual detail, occipital activation, and strengthened PPA–occipital coupling loaded negatively. This unsupervised structure reproduced the CTL > LB > CB gradient, showing that the perceptual–conceptual shift emerges not only in isolated measures but as a unified dimension that spans behavioral strategies, functional recruitment, and structural architecture. These results demonstrate that blindness reshapes mental time travel along a systematic representational axis rather than disrupting it.

### Limitations

Several limitations should be acknowledged. Our scoring schemes classify object-based descriptions as “visual,” which likely reflect conceptual rather than perceptual representations in congenitally blind participants, limiting the interpretability of visually coded details. The late-blind group showed substantial variability in age of onset and duration of blindness, constraining conclusions about developmental timing effects. Composite behavioral indices simplify interpretation but may obscure task-specific nuances, and neuroimaging results reflect group-level patterns that may mask individual variability in plasticity. These considerations qualify interpretation but do not alter the overall pattern of convergent behavioral and neural findings.

### Conclusion: Memory matters

Beyond its theoretical implications, this work speaks to the lived reality of people with blindness. Participants traveled from across Germany to take part in this study, underscoring how deeply memory matters in daily life. As one participant noted, “If I don’t remember that I put my cup of coffee in front of me, it is gone from my world.” Mental time travel is not an abstract cognitive function. It is the scaffold of identity, independence, and imagination. Our findings show that even without visual experience, the human episodic system preserves this capacity through flexible strategies, highlighting the resilience of memory and its central role in shaping identity and everyday cognition.

## Acknowledgements

We thank all participants for their time and commitment. We are especially grateful to the blind individuals who traveled long distances to Bonn to take part in this study. Their extraordinary effort made this research possible. We also thank Anke Rühling, Jennifer Schlee, and Yilmaz Sagik for their technical assistance during MRI scanning, and Patrick N. McCormick for helpful comments on the manuscript.

## Funding

This research was supported by the Hertie Network of Excellence in Clinical Neuroscience. Work in C.M.’s lab is further financed by internal research funding of the Faculty of Medicine (BONFOR), University Hospital Bonn, and by the Deutsche Forschungsgemeinschaft (DFG, German Research Foundation, MC 244/3-1 and project 493623632). Additional funding was provided by the Federal Ministry of Research, Technology and Space (BMFTR) under the funding code (FKZ): 01EO2107.

## Author Contributions

C.M. conceived and supervised the study, developed methodology, oversaw investigation and formal analyses, provided resources, and wrote the original draft. N.A.K. contributed to methodology, formal analysis, data curation, visualization, and writing the original draft. M.M. performed investigations, contributed to analyses, and assisted with manuscript editing. P.L. developed methods, conducted investigations and analyses (hippocampal volumes, fMRI, connectivity), and contributed to writing and review. J.T. contributed to methods and analyses (NLP and interview scoring). C.K. performed analyses of diffusion imaging. M.C. conducted behavioral interviews. A.E. contributed to methods and scoring analyses. S.L. assisted with methods, participant recruitment, and investigations. S.D. contributed to methods, investigations, data curation, and project administration. N.M. analyzed diffusion imaging data. E.G. conducted resting-state investigations. S.B. prepared technical setups for audio-MRI integration. T.S. contributed to MRI methodology and analyses. K.W. performed ophthalmological screening. B.W. assisted with participant recruitment and conceptual input. A.S. facilitated the research by offering space, assisting with participant recruitment, and contributing to manuscript drafting. All authors reviewed and approved the final manuscript.

## Competing Interests

The authors declare no competing interests.

## Supplementary Information

Supplementary Information includes:

- **Extended Methods:** Scoring schemes for autobiographical memory, scene construction, and episodic future thinking; interrater reliability; psychometric details (MoCA, BDI).
- **MRI Acquisition & Preprocessing:** Full parameters for task-based and resting-state fMRI, DTI, and structural imaging.
- **Statistical Models & Software**: ANCOVA/MANCOVA specifications, connectivity analyses, PCA procedures.
- **Additional Behavioral Analyses**: Perceptual–conceptual indices and sensory modality breakdowns by interview type.
- **Lexical Diversity Analysis**: Automated semantic metrics natural language processing and comparison with manual coding.
- **Neuroimaging Analyses**: Activation contrasts, connectivity profiles, VBM, hippocampal volumetry, and TBSS for white matter integrity.
- **Principal Component Analysis**: Integrated axis of mental time travel strategies combining behavioral and neural measures.
- **Additional Figures & Tables**: Extended behavioral results, semantic diversity plots, connectivity maps, and structural imaging contrasts.

## Data Availability Statement

All anonymized data and analysis code will be made available upon reasonable request to the corresponding author. Supplementary Information includes extended methods, scoring schemes, and additional analyses.

## Method summary

### Participants

We studied 81 adults (34 women; age range 18–81 years): 22 congenitally blind (CB), 22 late-blind (LB), and 37 sighted controls (CTL). Blindness was confirmed by ophthalmological examination; CB participants never had any visual experience, LB participants lost vision after age 2 (mean onset = 30.9 years, SD = 19.8). Groups were matched on sex and education; LB were older than CB and CTL (p = 0.04). Extensive supplementary methods provides details on causes of blindness, full demographic profiles, recruitment and ethical procedures, and group comparisons on general cognitive domains (MoCA, BDI). Ethical approval was obtained from the University Hospital Bonn ethics board. All participants gave written and oral informed consent and were reimbursed for participation.

### Design and Procedure

Participants completed behavioral interviews and MRI sessions. Not all participants completed all components (details in supplementary methods). Behavioral measures included:

- Mental imagery: Vividness of Visual Imagery Questionnaire (VVIQ)^23^ and Plymouth Sensory Imagery Questionnaire (PSIQ)^24^.
- Mental time travel (MTT): Three semi-structured interviews. i.e. Autobiographical Memory (AI)^23^, Scene Construction (SCI)^8^, and Episodic Future Thinking (EFT)^8^ adapted from established protocols.

Interviews were transcribed and scored for perceptual and conceptual details using validated schemes^8,25^. Interrater reliability and extended scoring procedures are described in supplementary methods.

### MRI Acquisition and Tasks

MRI data were collected on a 3T Siemens Skyra scanner (DZNE Bonn). Sessions included anatomical, diffusion tensor imaging (DTI), resting-state fMRI, and two task-based fMRI paradigms (AMI and SCI). Both tasks used auditory cues (realistic for AM, unrealistic for SC) followed by imagination phases (12 s) and vividness ratings. Math trials served as controls. Full acquisition parameters and preprocessing steps are provided in supplementary methods.

### Analysis

Behavioral data were analyzed using ANCOVA and MANCOVA models with age as covariate. Composite MTT scores contrasted perceptual and conceptual detail densities across interviews. Neuroimaging analyses included:

- Task-based activation contrasts (AM/SC vs. math) using SPM12 ttps://www.fil.ion.ucl.ac.uk/spm/).
- Resting-state and task-based connectivity analyses focused on the parahippocampal place area (PPA) using the conn toolbox v20.b (https://www.nitric.org/projects/conn/)
- White matter integrity assessed via tract-based spatial statistics (TBSS) on fractional anisotropy (FA) using FSL (https://fsl.fmrib.ox.ac.uk).
- Multimodal principal component analysis integrating behavioral and neural measures using SPSS.

Additional analyses (e.g., lexical diversity via large language models, hippocampal volumetry, voxel-based morphometry) and full statistical details are reported in supplementary methods.

## Supplementary Information

### Procedures

All experiments were conducted at the DZNE, Bonn, Germany. The study consisted of three main parts: (1) neuropsychological data collection, (2) the MRI block, and (3) an interview block. The order in which participants performed these blocks was dependent on the scanner schedule and the participants schedule (trains, hotels, etc.). The SCI was temporally conducted digitally.

### Participants

We included 81 adults (M = 54.14 years, SD = 1.86; range = 18–81; 34 women) across three groups: 22 congenitally blind (CB), 22 late-blind (LB), and 37 sighted controls (CTL). Extended demographic and cognitive characteristics are provided in Supplementary Table 1. Causes of blindness for CB and LB groups are listed in Supplementary Table 2.

LB participants lost vision between ages 2 and 76 (M = 30.86, SD = 19.75); blindness duration ranged from 4 to 58 years (M = 29.85, SD = 20.10). One CB participant was excluded due to incidental detection of a brain tumor during scanning. Blind participants were matched to CTL by age (±2 years), sex, and education. All participants gave written informed consent; the study was approved by the University Hospital Bonn ethics board. The study was pre-registered (osf.io./2zyev).

**Table S1.**
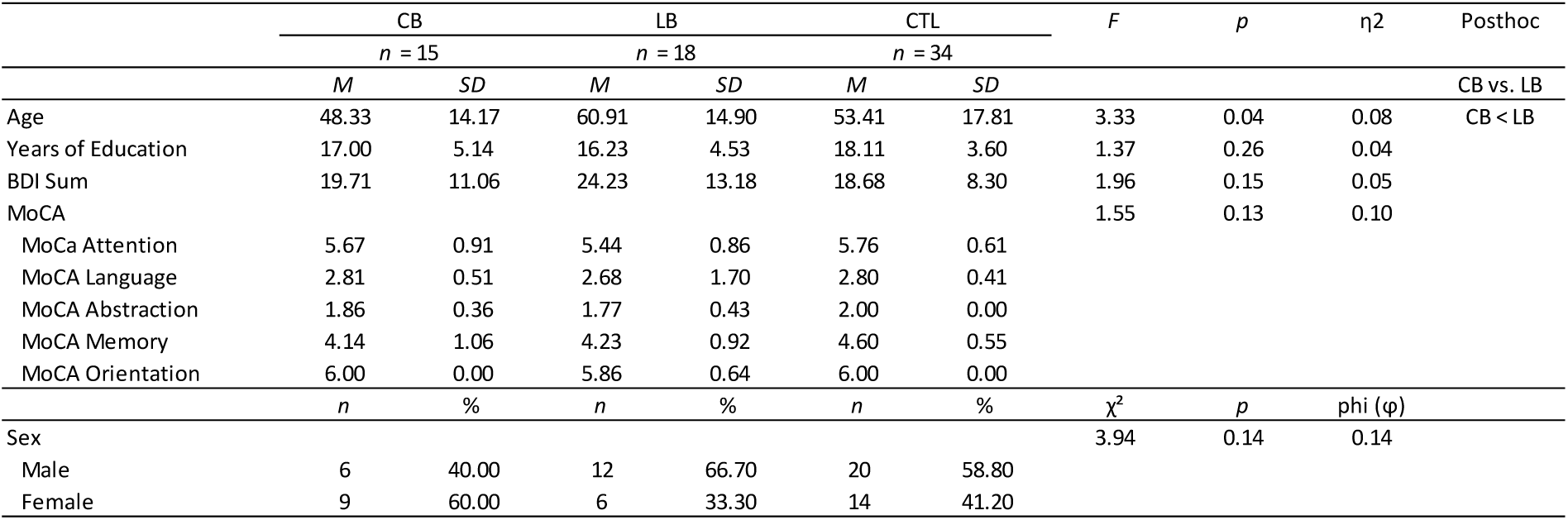
Demographical information. CB=congenital blind, LB=late blind, CTL=controls. BDI=Becks Depression Index, MoCA=Montreal cognitive assessment blind

| | CB | | LB | | CTL | | *F* | *p* | $\eta^2$ | Posthoc |
| --- | --- | --- | --- | --- | --- | --- | --- | --- | --- | --- |
|  | *n* = 15 |  | *n* = 18 |  | *n* = 34 |  |  |  |  |  |
|  | *M* | *SD* | *M* | *SD* | *M* | *SD* |  |  |  | CB vs. LB |
| Age | 48.33 | 14.17 | 60.91 | 14.90 | 53.41 | 17.81 | 3.33 | 0.04 | 0.08 | CB < LB |
| Years of Education | 17.00 | 5.14 | 16.23 | 4.53 | 18.11 | 3.60 | 1.37 | 0.26 | 0.04 |  |
| BDI Sum | 19.71 | 11.06 | 24.23 | 13.18 | 18.68 | 8.30 | 1.96 | 0.15 | 0.05 |  |
| MoCA |  |  |  |  |  |  | 1.55 | 0.13 | 0.10 |  |
| MoCa Attention | 5.67 | 0.91 | 5.44 | 0.86 | 5.76 | 0.61 |  |  |  |  |
| MoCA Language | 2.81 | 0.51 | 2.68 | 1.70 | 2.80 | 0.41 |  |  |  |  |
| MoCA Abstraction | 1.86 | 0.36 | 1.77 | 0.43 | 2.00 | 0.00 |  |  |  |  |
| MoCA Memory | 4.14 | 1.06 | 4.23 | 0.92 | 4.60 | 0.55 |  |  |  |  |
| MoCA Orientation | 6.00 | 0.00 | 5.86 | 0.64 | 6.00 | 0.00 |  |  |  |  |
| | *n* | % | *n* | % | *n* | % | $\chi^2$ | *p* | phi ($\phi$) | |
| Sex |  |  |  |  |  |  | 3.94 | 0.14 | 0.14 |  |
| Male | 6 | 40.00 | 12 | 66.70 | 20 | 58.80 |  |  |  |  |
| Female | 9 | 60.00 | 6 | 33.30 | 14 | 41.20 |  |  |  |  |

**Table S2.** Causes for blindness. CB=Congenital blind, LB=late blind.

| Cause of Blindness | CB<br>(N = 21) | LB<br>(N = 22) |
| --- | --- | --- |
| Retinis pigmentosa | 2 | 10 |
| Retinal degeneration | 4 | – |
| Retinal detachment | 2 | 4 |
| Glaucoma | 2 | 3 |
| Uveitis leading to Glaucoma | 1 | – |
| Peter-Plus-Syndrom leading to Glaucoma | – | 1 |
| Optic Neuropathy | 1 | – |
| Congenital Anirida | – | 1 |
| Leber Congenital Amaurosis (LCA) | 3 | – |
| Anolphthalmia | 1 | – |
| Accident | – | 3 |
| Unknown | 5 | – |

### Montreal Cognitive Assessment (MoCA)

Cognitive screening used a modified MoCA^1^ (max score = 22) excluding vision-dependent items^2^. A MANCOVA indicated no group differences across subscales, V = 0.20, F(10,142) = 1.58, p = .12.

### Beck’s Depression Inventory (BDI-V)

Depressive symptoms were assessed with the BDI-V^3,4^. Scores >35 indicate clinically relevant depression. Seven participants exceeded this cutoff (CB: n = 2; LB: n = 4; CTL: n = 1).

### Mental Imagery questionnaires

#### Description

Mental imagery was assessed using German versions of two validated self-report instruments: (1) Vividness of Visual Imagery Questionnaire (VVIQ)^5^ and (2) Plymouth Sensory Imagery Questionnaire (PSIQ) short form^6^. The VVIQ comprises 16 items prompting participants to imagine four scenes (e.g., a rising sun) and rate image vividness on 5-point Likert scales (1 = no image, 5 = perfectly clear and vivid). The PSIQ evaluates imagery across seven modalities (vision, sound, smell, taste, touch, bodily sensation, emotional feeling) using three items per modality. Vividness ratings are provided on 10-point scales (0 = no image, 10 = as vivid as real life). Both questionnaires have demonstrated good psychometric properties in German samples^7^.

#### Statistical Results

An ANCOVA controlling for age revealed significant group differences in VVIQ scores, F(2,75) = 29.81, p < .001, η² = .44. Post-hoc tests (Bonferroni) showed CB < CTL (mean difference = 24.93, SE = 3.40, p < .001) and LB < CTL (mean difference = 24.23, SE = 3.89, p < .001); CB and LB did not differ significantly.

For PSIQ, overall scores did not differ across groups, F(2,74) = 0.39, p = .68, η² = .01, but a MANCOVA on modality-specific subscores revealed a significant group effect, Pillai’s Trace = .65, F(14,136) = 4.70, p < .001. Follow-up ANCOVAs indicated a pronounced effect on the vision subscale, F(2,73) = 37.52, p < .001, η² = .51, with CB scoring significantly lower than LB and CTL (p’s < .001). To test whether CB participants scored above the aphantasia cutoff (VVIQ = 32), a one-sample t-test was conducted. CB scores were significantly higher than the cutoff, t(21) = 3.1, p = .007, confirming that congenitally blind individuals do not meet criteria for aphantasia despite reduced visual imagery vividness.

### Narrative data

#### 6.1 Autobiographical Memory interview

##### Description

Autobiographical performance was assessed using semi-structured Autobiographical Memory Interviews (AMI^8^). Participants recalled events from five predefined life periods:

- early childhood (≤11 years),
- adolescence (11–17 years),
- early adulthood (18–35 years),
- middle age (35–55 years), and
- the previous year. Each interview comprised:
- Free recall phase: Participants described the event until reaching a natural endpoint; general probes were used if detail was sparse.
- Structured probe phase: Interviewers asked specific questions about each event.
- Subjective ratings: Emotionality, importance, and rehearsal frequency on 6-point Likert scales.

Two independent raters transcribed and scored interviews using Levine’s scheme, dividing details into:

- Internal details (episodic): event, time, place, perception, emotion/thought.
- External details: semantic info, repetitions, unrelated events.

**Methodological extensions**:

- Expanded perceptual coding: Sensory modalities scored separately (visual, auditory, tactile, gustatory, olfactory).
- Additional categories: physical sensations, body position, spatial body position, perceived duration.
- Qualitative ratings: Episodic richness, time integration, temporal extension, coherence.

##### Debriefing

Ten open-ended questions explored memory strategies, spatial orientation, imagination, and social integration.

#### Statistical results

##### Interrater Reliability

Composite scores for internal and external details showed excellent reliability (ICC, one-way random effects): internal = .95; external = .90.

##### Verbosity and Detail Counts

To control for individual differences in verbosity, detail counts were converted to density scores (details per word), following recommendations that verbosity-adjusted metrics provide more stable and valid measures of episodic performance^9,10^. No group differences emerged for total word count, F(2,75) = 0.37, p = .69, η² = .01, or code count, F(2,75) = 0.14, p = .87, η² = .00. Age had no effect (p’s > .38).

##### Density Scores

Standardizing detail counts by word count revealed a significant group effect for internal detail density, F(2,75) = 6.37, p = .003, η² = .15. CB scored lower than CTL (mean difference = 0.003, SE = 0.001, p = .002). External detail density did not differ, F(2,75) = 1.70, p = .20.

##### Internal Densities Subcategories

MANCOVA on internal detail densities (event, time, place, perception, emotion/thought) showed a significant group effect, Pillai’s Trace = .34, F(8,146) = 3.67, p < .001, η² = .17. CB reported more time details than CTL but fewer perceptual details than both other groups.

##### Perceptual Densities Subcategories

MANCOVA on perceptual detail densities (vision, audition, touch, gustation, olfaction, body position, spatial body position, physical sensation, duration) indicated a strong group effect, Pillai’s Trace = .74, F(16,138) = 5.10, p < .001, η² = .37.

**Follow-up ANCOVAs**:

- Vision: F(2,75) = 39.96, p < .001, η² = .52
- Audition: F(2,75) = 17.79, p < .001, η² = .32
- Tactile: F(2,75) = 5.37, p = .007, η² = .13
- Body position: F(2,75) = 4.58, p = .013, η² = .11
- Duration: F(2,75) = 8.30, p < .001, η² = .18

CTL showed highest vision density; LB intermediate; CB lowest. CB exceeded CTL in tactile and duration details; both blind groups exceeded CTL in auditory details. Within-group proportions: CTL narratives were ∼50% visual; CB narratives included ∼25% visual and ∼20% auditory details (Fig. 1). CB participants reported significantly more visual than auditory details, paired t-test: t(20) = 2.584, p = .018 (two-tailed).

##### Subjective Ratings

Group differences emerged for ability to visualize episodes, F(2,74) = 6.12, p < .01, η² = .14 (CB < LB and CTL), and for present importance ratings, F(2,74) = 6.48, p < .01, η² = .15 (CTL < CB and LB). Past importance ratings were higher in CB than LB (p = .04). No differences for rehearsal frequency or emotional impact (p’s > .24).

##### Subjective Episodic Richness and Re-experiencing

No group differences for episodic richness, F(2,75) = 0.58, p = .56, or composite re-experiencing score, F(2,75) = 0.39, p = .68. MANCOVA on ratings (time, place, perception, thought/emotion) revealed a significant group effect, Pillai’s Trace = .31, F(8,146) = 3.35, p < .01, η² = .16. LB scored higher on time ratings than CTL; CB scored lower on perception ratings than CTL.

##### Debriefing

All groups rated memory retrieval as easy (means 1.55–2.38 on a 6-point scale). CTL found retrieval slightly more difficult than LB, F(2,73) = 4.34, p = .02, η² = .11. None reported loneliness.

**Supplementary Fig. S1.**
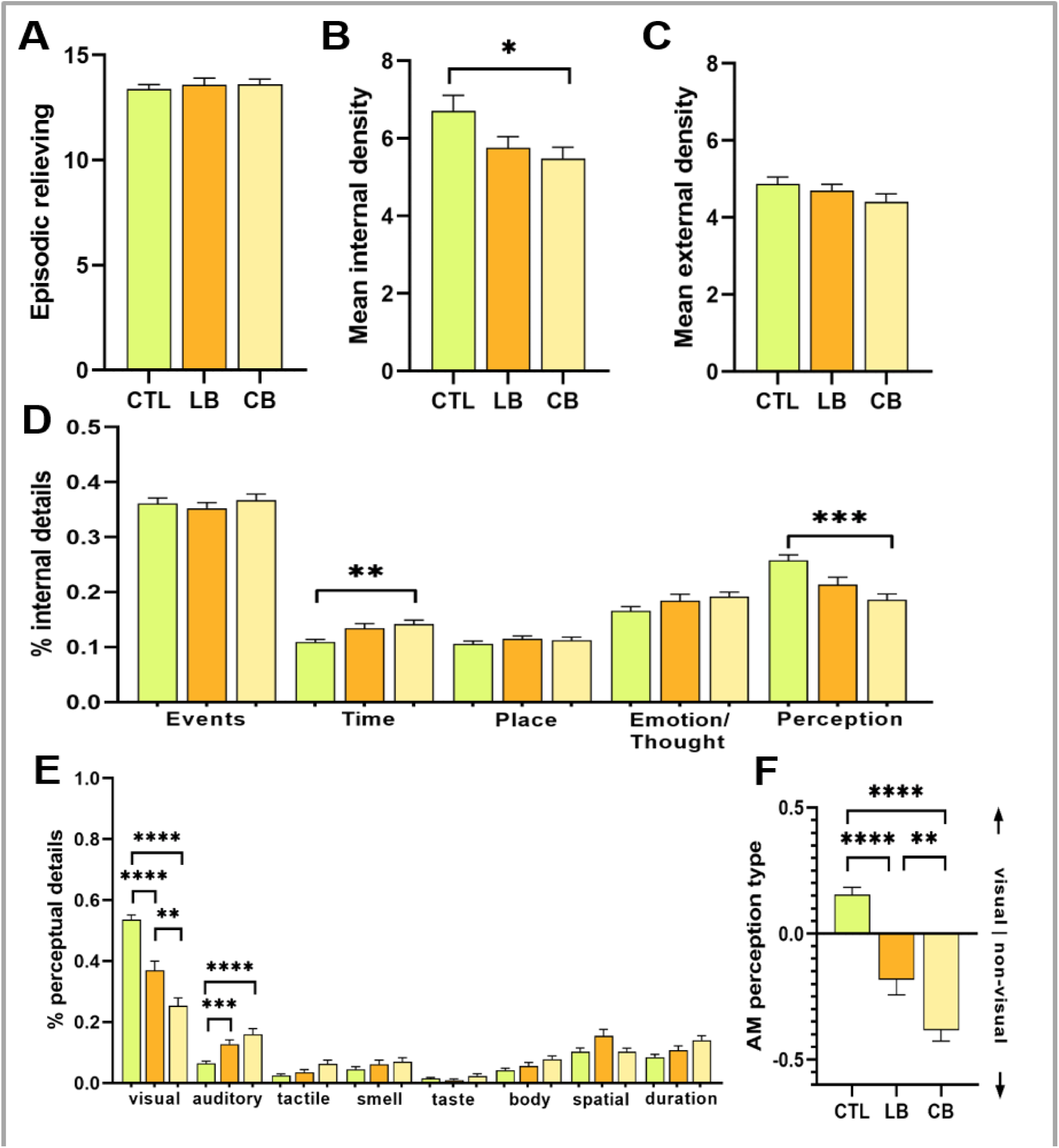
Autobiographical memory profiles differ by visual experience. (a) Subjective ratings of episodic reliving did not differ significantly between groups (CTL, LB, CB). (b) Internal detail density was highest in sighted controls (CTL), intermediate in late-blind (LB), and lowest in congenitally blind (CB) (p < 0.05). (c) External detail density showed no significant group differences. (d) Composition of internal details revealed modality-specific adaptations: CB participants reported fewer perceptual details and more conceptual details (emotion/thought) compared to CTL, with LB showing an intermediate profile (p < 0.01 for time; *p < 0.001 for perception). (e) Sensory breakdown of perceptual details indicated visual dominance in CTL, mixed profiles in LB, and non-visual reliance (auditory, tactile) in CB (**p < 0.0001 for visual vs. auditory/tactile contrasts). (f) perceptual–conceptual index for autobiographical memory confirmed perceptual dominance in CTL and conceptual dominance in CB, with LB intermediate (**p < 0.0001). Bars represent means ± SEM.

#### 6.2 Scene construction and episodic future thinking

##### Description

Scene construction and episodic future thinking were assessed using the SCI protocol^11^. Participants imagined and described 10 scenarios:

- Atemporal scene construction: first seven scenarios (e.g., “Imagine you are sitting in an elegant seminar room at a large company”).
- Episodic future thinking: last three scenarios (e.g., “Imagine you are attending a future celebration”).

After each scenario, participants rated sense of presence, perceived salience, and spatial coherence. Sighted participants completed questionnaires independently; blind participants received verbal instructions.

##### Scoring

Two independent raters coded transcripts for: Entities present, Thoughts/emotion/action, Spatial references, Sensory descriptions (vision, audition, tactile, olfactory, gustatory). Two independent raters provided quality judgments from 0 to 10. The main outcome was the experiential index, combining subjective ratings, coded details, and qualitative judgments. Separate indices were computed for atemporal scene construction and episodic future thinking.

##### Debriefing

Participants answered open-ended questions on how they imagine their environment and orient themselves at home and in unfamiliar places.

#### Statistical results for atemporal scene construction

##### Word Counts and Verbosity (SC)

For atemporal scene construction (SC), no group differences were found in word count, F(2,63) = 0.78, p = .46, η² = .02, or total code count, F(2,63) = 1.34, p = .27, η² = .04. Word count correlated strongly with detail count across the sample, r(68) = .78, p < .001. Verbosity-adjusted content density showed a significant group effect, F(2,63) = 3.50, p = .036, η² = .10. Pairwise comparisons indicated a trend for CB to score lower than LB (mean difference = –.04, SE = .01, p = .059) and CTL (mean difference = –.03, SE = .01, p = .059).

##### Content Detail Ratios (SC)

MANCOVA on ratios of spatial references, sensory descriptions, and thoughts/emotions/actions revealed a significant group effect, Pillai’s Trace = .21, F(6,124) = 2.37, p = .034, η² = .10, and an age effect, Pillai’s Trace = .48, F(3,61) = 18.52, p < .001, η² = .48.

**Follow-up ANCOVAs (SC)**:

- Thoughts/emotions/actions: F(2,63) = 5.86, p = .005, η² = .16 (CB > LB and CTL)
- Spatial references: F(2,63) = 4.37, p = .017, η² = .12 (CTL > CB)

##### Perceptual Detail Ratios (SC)

MANCOVA on perceptual modalities (vision, audition, touch, smell, taste) showed a significant group effect, Pillai’s Trace = .55, F(8,122) = 5.77, p < .001, η² = .27, and an age effect, Pillai’s Trace = .21, F(4,60) = 3.06, p = .023, η² = .17.

**Follow-up ANCOVAs (SC)**:

- Vision: F(2,63) = 32.36, p < .001, η² = .51
- Audition: F(2,63) = 22.59, p < .001, η² = .42
- Touch: F(2,63) = 13.46, p < .001, η² = .30

CTL reported the highest proportion of visual details; CB the lowest. Blind groups reported more auditory and tactile details than CTL.

**Supplementary Fig. S2.**
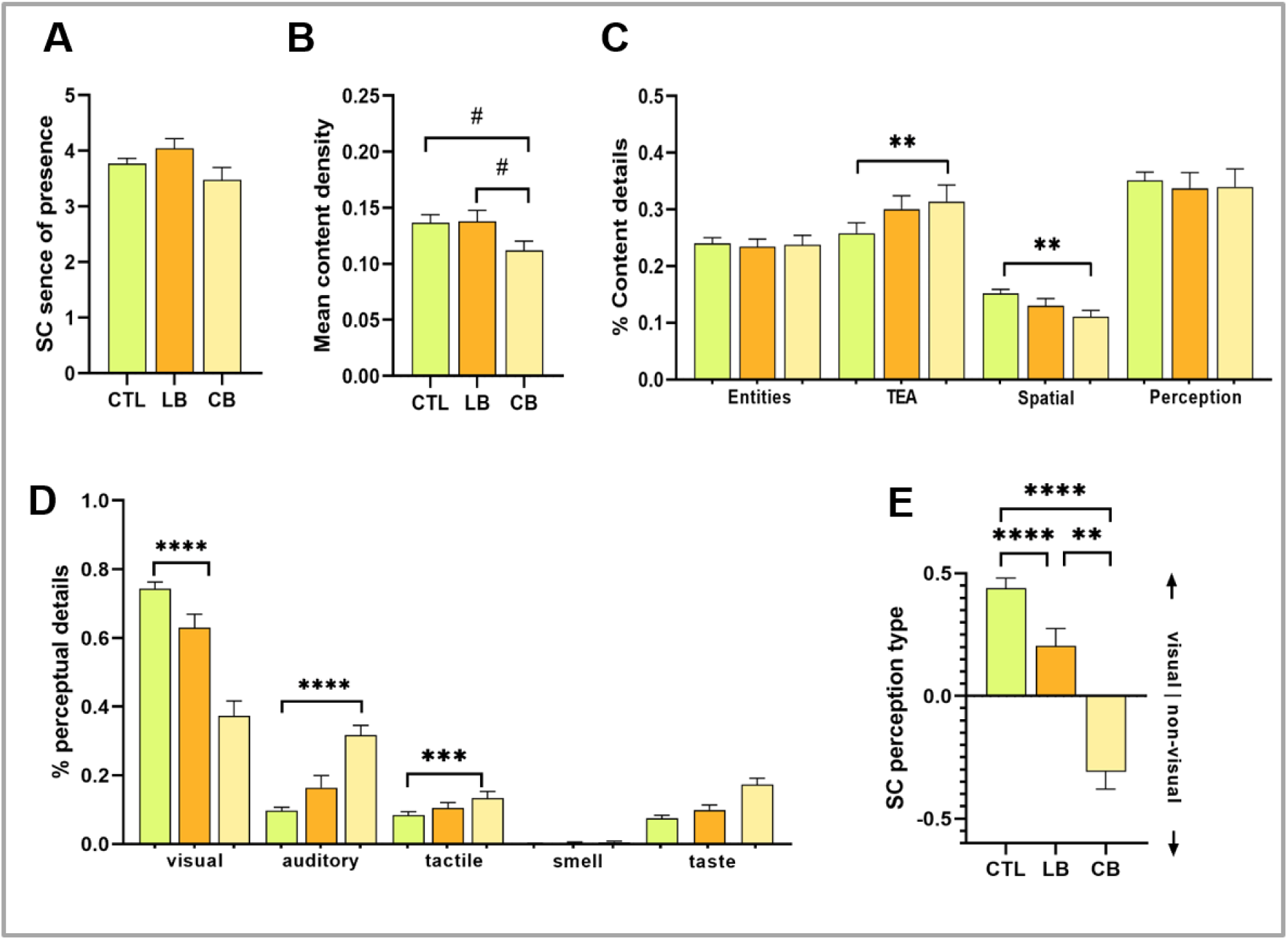
Scene construction profiles differ by visual experience. (a) Subjective ratings of sense of presence did not differ significantly between groups (CTL, LB, CB). (b) Mean content density was highest in sighted controls (CTL) and lowest in congenitally blind participants (CB), with LB intermediate (trend level, p’s < 0.07). (c) Composition of content details revealed group differences: CTL produced more spatial references, whereas CB emphasized thoughts/emotions/actions (TEA, p < 0.01 for TEA and spatial categories). (d) Sensory breakdown of perceptual details indicated visual dominance in CTL, mixed profiles in LB, and non-visual reliance (auditory, tactile) in CB (**p < 0.0001 for visual vs. auditory/tactile contrasts; *p < 0.001 for tactile). (e) Perceptual–conceptual index for scene construction confirmed perceptual dominance in CTL and conceptual dominance in CB, with LB intermediate (**p < 0.0001). Bars represent means ± SEM.

#### Statistical results for episodic future thinking

##### Word Counts and Verbosity (EFT)

No group differences in word count, F(2,63) = 1.20, p = .31, η² = .04, or total content count, F(2,63) = 0.75, p = .48, η² = .02. Verbosity-adjusted scores did not differ, F(2,63) = 0.76, p = .47, η² = .02. Word count correlated with detail count, r(68) = .72, p < .001.

##### Content Detail Ratios (EFT)

MANCOVA on ratios of spatial references, entities present, thoughts/emotions/actions, and sensory descriptions showed a trend for group effect, Pillai’s Trace = .18, F(6,124) = 2.37, p = .07, η² = .10.

##### Perceptual Detail Ratios (EFT)

MANCOVA on perceptual modalities revealed a significant group effect, Pillai’s Trace = .35, F(8,116) = 3.05, p = .004, η² = .17. Significant effects for vision and audition; trend for tactile. CB reported fewer visual but more auditory details than CTL.

##### Debriefing

Blind participants emphasized auditory and tactile modalities for imagining and orientation, followed by olfactory cues. Both groups reported using tools (guide dogs, canes, digital navigation) and environmental markers outside the home; at home, orientation relied on memory and tactile markers. Some LB participants described imagining scenarios visually “as if sighted,” but their VVIQ scores did not differ from other LB participants.

**Supplementary Fig. S3.**
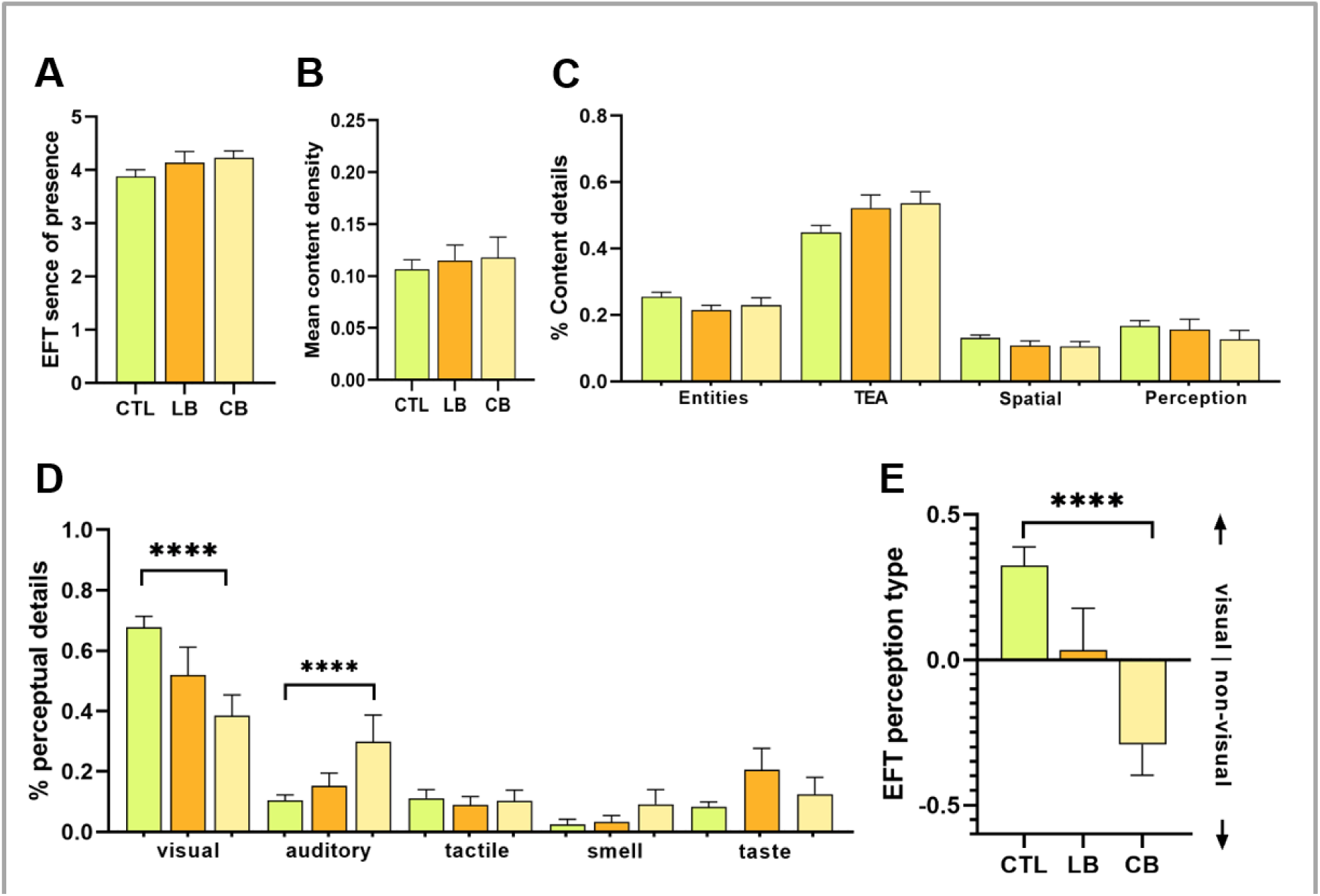
Episodic future thinking profiles differ by visual experience. (a) Subjective ratings of sense of presence did not differ significantly between groups. (b) Mean content density showed no significant group differences. (c) Composition of content details revealed similar patterns across groups, with CB emphasizing thoughts/emotions and CTL producing more spatial references. (d) Sensory breakdown of perceptual details indicated strong visual dominance in CTL, mixed profiles in LB, and non-visual reliance in CB (**p < 0.0001 for visual vs. auditory contrasts). (e) Perceptual–conceptual index for episodic future thinking confirmed perceptual dominance in CTL and conceptual dominance in CB, with LB intermediate (**p < 0.0001). Bars represent means ± SEM.

#### 6.3 Mental Time Travel (MTT) Score

##### Description

To capture cross-modal strategies, we computed a Mental Time Travel (MTT) score ranging from –1 to +1, reflecting the balance between conceptualization (negative values) and perceptual detail (positive values). For AI, conceptual scores were based on emotion/thought detail densities. For SC and EFT, conceptual scores were based on thoughts/emotion/action detail densities. Perceptual scores combined perceptual detail densities and spatial references (included for consistency with AMI, where perception includes spatial positioning).

The MTT score was calculated as: MTT = (Perceptual Density) − (Conceptual Density)

Negative values indicate conceptual dominance; positive values indicate perceptual dominance. Finally, the mean of the three task-specific scores (AMI, SC, EFT) yielded a cross-modal conceptual–perceptual index.

##### Statistical results

Significant group effects were observed for the composite MTT score across all tasks:

- AI: F(2,75) = 10.63, p < .001, η² = .22
- SC: F(2,63) = 5.92, p = .004, η² = .16
- EFT: F(2,63) = 4.00, p = .02, η² = .11

Post-hoc comparisons (Bonferroni-adjusted):

- CB scored significantly lower than CTL for AM (M = –.11, SE = .02, p < .001) and EFT (mean difference = –0.21, SE = .07, p = .02).
- For SC, CB scored lower than LB (mean difference = –0.15, SE = .06, p = .043) and CTL (mean difference = –0.17, SE = .05, p = .003).
- No significant differences between LB and CTL for any task.

Cross-modal index. Groups differed significantly on the integrated conceptual–perceptual index, F(2,63) = 8.56, p < .001, η² = .24 (Fig. 2).

#### Semantic diversity: Natural language processing

##### Description

Interview transcripts from AMI, SC, and EFT were preprocessed (lowercasing, whitespace and non-alphanumeric removal, German stop-word filtering, stemming via NLTK^12^) and tokenized at the word level^13^. For each group, word frequencies were computed and converted to ranks (rank 1 = most frequent). Using ranks instead of raw counts mitigates verbosity differences. ). Higher diversity indicates idiosyncratic, richly varied descriptions, whereas lower diversity reflects semantically closer constructs and reliance on stereotypical templates.

Lexical diversity was indexed by median word rank per subject across all tokens, averaged across AMI, SC, and EFT. Higher ranks indicate greater diversity (more unique words), whereas lower ranks reflect reduced diversity and more semanticized, template-like language. Group differences were tested using one-way ANOVA with Tukey’s HSD post hoc comparisons.

##### Statistical Results

CTL showed the highest ranks (M = 221.48, SD = 13.33), followed by LB (M = 166.39, SD = 11.42) and CB (M = 133.68, SD = 10.72). ANOVA confirmed a significant group effect, F(2,76) = 52.92, p < .001 (4.01e–15), partial η² = .58.

Post hoc tests indicated CTL > LB > CB:

- CTL vs. LB: p < .001, d = 1.58
- CTL vs. CB: p < .001, d = 2.55
- LB vs. CB: p = .004, d = 1.32

#### Neuroimaging

### 8.1 Scanner and hardware

Data were acquired on a 3T Siemens Skyra scanner (Siemens Healthineers, Erlangen, Germany) at the German Center for Neurodegenerative Diseases (DZNE), Bonn, using a 32-channel head coil.

### 8.2 Functional MRI

#### 8.2.1 Functional MRI acquisition

Task-based fMRI for autobiographical memory and scene construction employed a gradient-echo EPI sequence with TR = 2000 ms, TE = 30 ms, and a voxel size of 3.5 mm isotropic. The acquisition matrix was 64 × 64 with 39 slices oriented oblique-axially along the AC–PC line. The bandwidth was 2112 Hz/pixel, and each session comprised 460 volumes with a total acquisition time of approximately 15 minutes. Resting-state fMRI used the same EPI parameters as the task-based scans, with 190 volumes acquired over approximately 7 minutes. Participants were instructed to keep their eyes closed and think of nothing during this scan.

#### 8.2.2 Functional MRI stimulus delivery and timing

##### Stimulus Delivery and Timing

The paradigm was adapted from previous research^14–16^. Before scanning, participants completed practice trials to familiarize themselves with the tasks. During the experiment, auditory cues were presented via MRI-compatible headphones, and responses were recorded using button presses. The autobiographical memory (AM) task comprised 30 randomized trials (15 AM and 15 math control trials). For AM trials, participants heard an event cue (e.g., “a kiss,” “visiting the zoo”) and retrieved a specific autobiographical memory. Once a memory was identified, they pressed a button and spent 12 seconds mentally reliving the event. Math trials involved simple calculations (e.g., 19 + 4); after solving, participants pressed a button and continued adding three to the result until the 12-second interval ended. After each trial, participants rated vividness for AM trials or difficulty for math trials. Inter-stimulus intervals were jittered between 1 and 4 seconds.

The scene construction (SC) task followed the same timing structure but used unrealistic auditory cues (e.g., “meeting a fairy”) to ensure novelty and avoid overlap with autobiographical events. It comprised a total of 40 randomized (20 SC and 20 math trials). Participants imagined each scenario in detail and rated vividness after each trial. Math trials served as controls and were rated for difficulty.

#### 8.2.3 Functional MRI preprocessing

Functional MRI data from the autobiographical memory (AM) and scene construction (SC) tasks were preprocessed using SPM12 (Wellcome Trust Centre for Neuroimaging, London, UK) implemented in MATLAB R2022a. The pipeline began with reorientation of all images to the AC-PC line, followed by realignment of all functional volumes to the first image to correct for head motion. Geometric distortions were minimized through unwarping using voxel displacement maps derived from field maps. Functional scans were co-registered to the bias-corrected structural T1-weighted image and normalized to MNI152 space using the unified segmentation approach. Finally, data were smoothed with an 8 mm FWHM Gaussian kernel.

##### Task-based fMRI analyses

For activation analyses, first-level models included separate regressors for autobiographical memory (AM) and scene construction (SC) trials, each contrasted against math control trials, with six motion parameters entered as nuisance covariates. The main contrasts of interest were AM > Math and SC > Math. To increase power and capture shared processes, we additionally computed a unified contrast combining both tasks (AM + SC > AM-Math + SC-Math) using the SPM concatenation pipeline. Statistical maps were thresholded at voxel-wise p < 0.05 FDR corrected and exceeding a cluster size of 50 adjacent voxels. At the second level, we implemented an ANCOVA with age as a covariate of no interest to compare the three groups (CTL, LB, CB), testing for main effects and group × task interactions. Regions showing significant interaction effects were subjected to follow-up ANOVAs on extracted beta estimates to characterize group differences.

##### Resting state and task-based functional connectivity analysis

Functional connectivity was examined using the CONN toolbox v20.b (https://www.nitric.org/projects/conn/) for both resting-state and task-based fMRI data. For connectivity analyses, we focused on ROI-to-ROI correlations within the episodic network, using seeds in the parahippocampal place area (PPA) and posterior cingulate cortex (PCC). ROIs were defined as 10 mm spheres centered on peak coordinates identified in the task-based interaction analysis (including rOCC, lOCC, fusiform gyrus, and PCC). Connectivity maps were thresholded at FDR-corrected p < 0.05 with a minimum cluster size of 50 contiguous voxels.

#### 8.2.4 Task-based fMRI behavioral results

During scanning, all groups performed the autobiographical memory (AM) and scene construction (SC) tasks with high compliance and comparable behavioral profiles. For the AM task, vividness ratings did not differ significantly between groups (Pillai’s Trace = 0.01, F(2,65) = 0.32, p = .73). Sighted controls rated 81% of AM trials as vivid, congenitally blind participants 80%, and late-blind participants 71%. Reaction times showed no interaction between trial type and group (F(2,65) = 1.12, p = .50), and missing responses were rare, averaging 10.8% for AM trials and 8.8% for math trials without group differences.

For the SC task, vividness ratings were generally lower but again showed no significant group effect (Pillai’s Trace = .026, F(2,64) = 0.64, p = .53). Sighted controls rated 73.4% of SC trials as vivid, compared to 59% in the congenitally blind group and 57% in the late-blind group. Math trials were rated as easy in over 90% of cases by controls and congenitally blind participants, and in 84% by late-blind participants. Reaction times did not differ across groups (F(2,64) = 1.55, p = .22), and missing data were minimal (9.5% for SC trials, 7.7% for math trials) with no group differences.

Overall, these findings indicate that task compliance was high across all groups, with vividness ratings for AM trials near ceiling and SC trials moderately lower, but without significant behavioral differences.

#### 8.2.5 Task-based MRI: Activation results

Task-based fMRI during autobiographical memory (AM, Fig. S4) and scene construction (SC, Fig. S5) revealed a common episodic network across all groups, including bilateral hippocampus, parahippocampal cortex, posterior cingulate cortex (PCC), lateral parietal cortex, and ventromedial prefrontal cortex (vmPFC).

In both tasks, group-level interactions indicated similar modality-specific adaptations. Sighted participants exhibited stronger activation in regions supporting visuospatial scene construction, notably the parahippocampal place related-area (PPA, MNI 28 -32 -22) and right fusiform gyrus (MNI 26 -62 -12). To label scene-selective peaks, we used the anatomical definition of the PPA as lying at the junction of the collateral and anterior lingual sulci, following Weiner et al. (2018). Coordinates falling within this anatomical zone were referred to as PPA-related.

In contrast, congenitally blind participants showed greater engagement of conceptual hubs, including PCC (MNI -2 -64 42) and bilateral lateral occipital cortex (right OCC, MNI 14 -64 0 and left OCC, MNI -12 -68 2), regions typically associated with visual processing but repurposed here through cross-modal plasticity (ANCOVA with age as covariate). Late-blind participants displayed intermediate profiles, consistent with partial retention of visual imagery.

**Supplementary Fig. S4.**
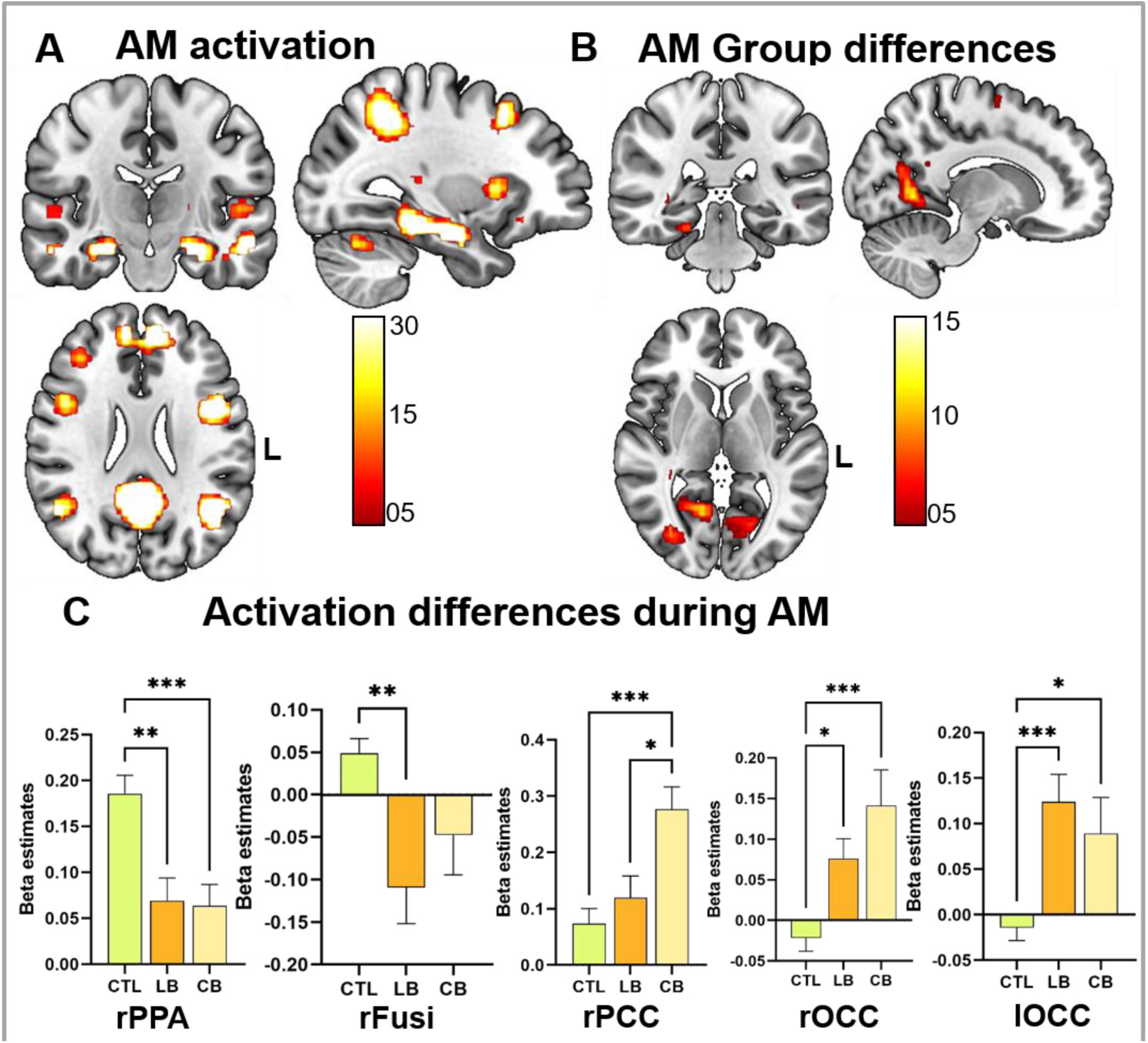
Neural architecture during autobiographical memory (AM) and group-specific adaptations. (a) Task-based fMRI during autobiographical memory revealed a common episodic network across all groups, including bilateral hippocampus, parahippocampal cortex, posterior cingulate cortex (PCC), lateral parietal cortex, and ventromedial prefrontal cortex (vmPFC). (b) Group differences emerged in regions supporting mental time travel: sighted participants showed stronger activation in the parahippocampal place area (PPA) and fusiform gyrus, whereas congenitally blind participants exhibited greater engagement of PCC and lateral occipital cortex (LOC). (c) Beta estimates from regions showing significant group differences illustrate effect magnitude and direction. Sighted controls (CTL) exhibited higher activation in PPA and fusiform gyrus, while congenitally blind participants (CB) showed increased activation in PCC and occipital regions (rOCC, lOCC). Late-blind participants (LB) displayed intermediate profiles. Bars represent means ± SEM; significance markers indicate ANCOVA results (p < 0.01**, \***p < 0.001**, \*\***p < 0.0001).

**Supplementary Fig. S5.**
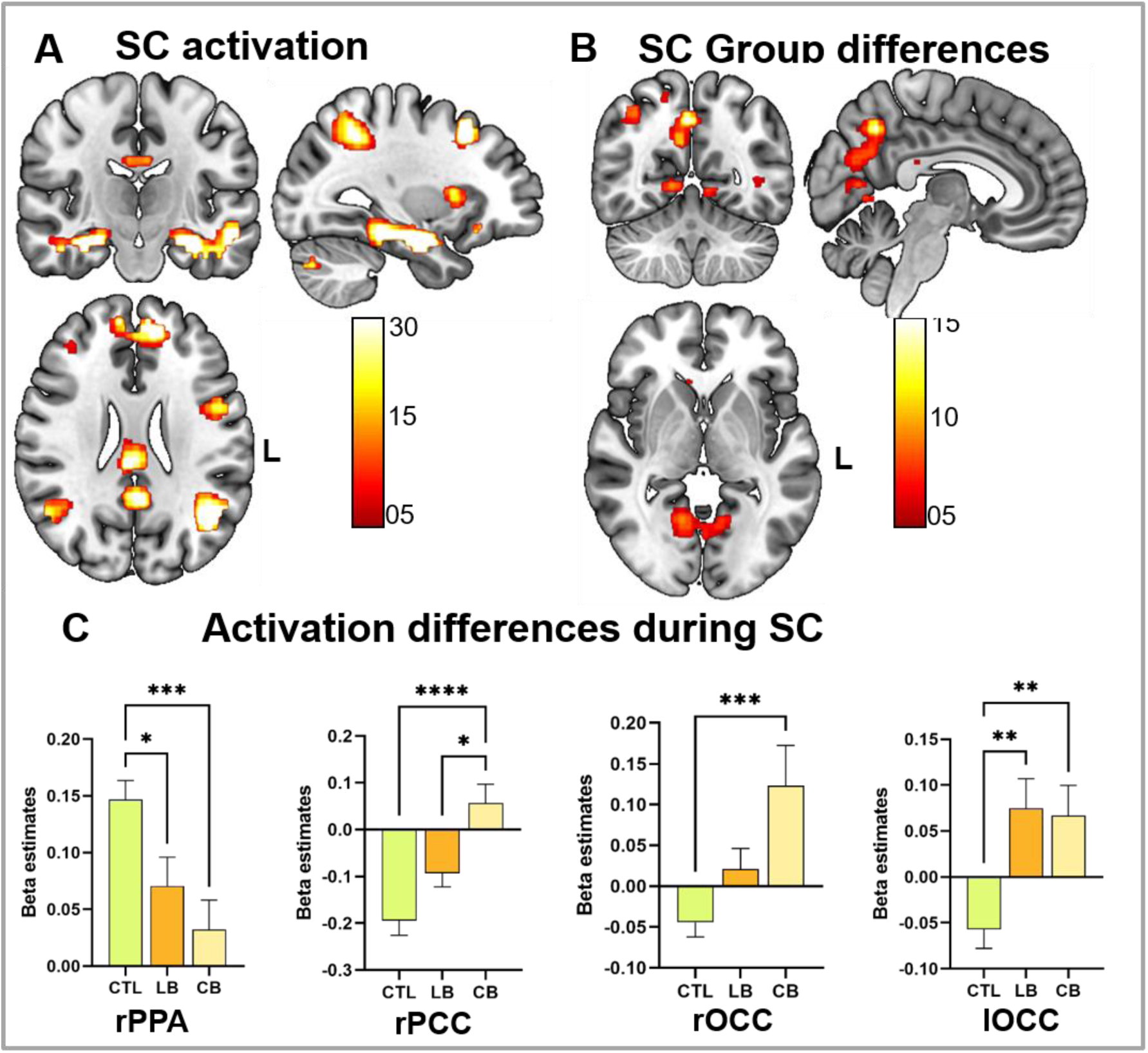
Neural architecture during scene construction (SC) and group-specific adaptations. (a) Task-based fMRI during scene construction revealed a common episodic network across all groups, including bilateral hippocampus, parahippocampal cortex, PCC, lateral parietal cortex, and vmPFC. (b) Group differences mirrored those observed for autobiographical memory: sighted participants showed stronger activation in PPA, whereas congenitally blind participants exhibited greater engagement of PCC and occipital cortex (LOC). (c) Beta estimates from regions showing significant group differences illustrate effect magnitude and direction. Sighted controls (CTL) exhibited higher activation in PPA, while congenitally blind participants (CB) showed increased activation in PCC and occipital regions (rOCC, lOCC). Late-blind participants (LB) displayed intermediate profiles. Bars represent means ± SEM; significance markers indicate ANCOVA results (p < 0.01, *p < 0.001, **p < 0.0001).

#### 8.2.6 Brain-Behaviour interactions

##### Description

To examine the neural basis of perceptual versus conceptual strategies in mental time travel, we correlated task-based fMRI activation in key regions with behavioral indices. Specifically, beta estimates from brain regions that showed marked group differences, namely the parahippocampal place area (PPA), fusiform gyrus (Fusi), posterior cingulate cortex (PCC), and bilateral occipital cortex (rOCC, lOCC) were extracted for the concatenated autobiographical memory and scene construction tasks. In addition, we extracted pre-specified regions of interest, such as the right and left hippocampus (MNI: 26 -18 -20 and -24 -18 -20) and the vmPFC (MNI 0 48 -14). Lastly, we extracted beta estimates form the right dorsolateral PFC (MNI 42 22 24) as a control region, for which we expected no correlation with our behavioural measures. Behavioral predictors included perceptual detail density and conceptual detail density. Correlations were computed across all participants and within groups using Pearson’s r, with significance assessed via two-tailed tests.

##### Statistical results

rPPA activation correlated positively with MTT visual detail density (r = 0.45, p < 0.001) and negatively with non-visual detail density (r = -0.35, p = 0.004). rFusiform activation a similar pattern but with weaker associations (visual details: r = 0.17, p = 0.18, non-visual details r = - 0.13, p = 0.33). We found the same pattern for regions typically associated with visual imagery, such as the right (visual details: r = 0.34, p = 0.005, non-visual details r = -0.29, p = 0.02) and left hippocampus (visual details: r = 0.32, p = 0.007, non-visual details r = -0.28, p = 0.01), as well as the vmPFC (visual details: r = 0.28, p = 0.02, non-visual details r = -0.24, p = 0.04).

rOCC activation showed the opposite pattern: positive correlation with non-visual detail density (r = 0.35, p = 0.004) and negative correlation with visual detail density (r = -0.39, p = 0.002). lOCC exhibited a similar pattern to that of the rOCC (visual details: r = -0.33, p = 0.008, non-visual details r = 0.22, p = 0.08). PCC exhibited a similar pattern to that of the right and left OCC (visual details: r = -0.34, p = 0.005, non-visual details r = 0.32, p = 0.009).

Lastly, we did not find any significant correlations in our control brain region, the dorsolateral PFC (visual details: r = -0.12, p = 0.17, non-visual details r = 0.16, p = 0.34).

**Supplementary Fig. S6.**
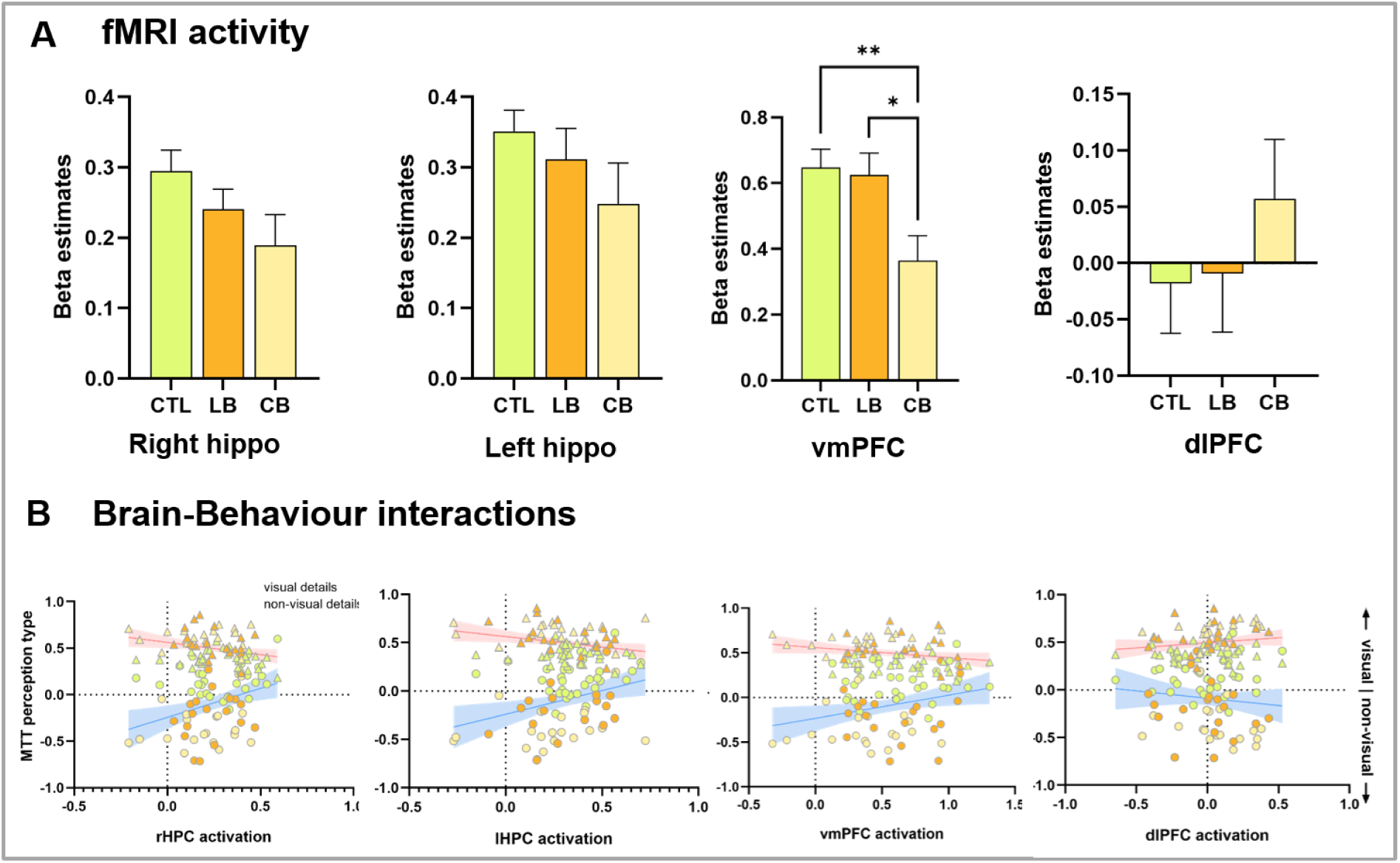
Brain–behavior interactions within key regions of the episodic network. (a) Extracted beta estimates from the right and left hippocampus, ventromedial prefrontal cortex (vmPFC), and a control region (dorsolateral prefrontal cortex, dlPFC) during autobiographical memory and scene construction tasks. Bars represent mean activation (± SEM) across groups. (b) Scatterplots showing correlations between beta estimates and behavioral indices: visual detail density (blue) and non-visual detail density (red) derived from interview scoring. Both hippocampi and vmPFC exhibit positive associations with visual details and negative associations with non-visual details, whereas dlPFC shows no significant correlations. Regression lines indicate best-fit trends; shaded areas represent 95% confidence intervals.

#### 8.2.7 Task-based fMRI: Functional connectivity results

Task-based connectivity analyses, separately conducted during autobiographical memory and scene construction revealed group-specific adaptations within the episodic network. Compared to sighted controls, congenitally blind participants showed significantly stronger functional coupling between the parahippocampal place area (PPA) and bilateral occipital cortex, as well as enhanced connectivity with fusiform gyrus. Late-blind participants exhibited intermediate connectivity profiles, consistent with partial retention of visual strategies.

**Figure S7.**
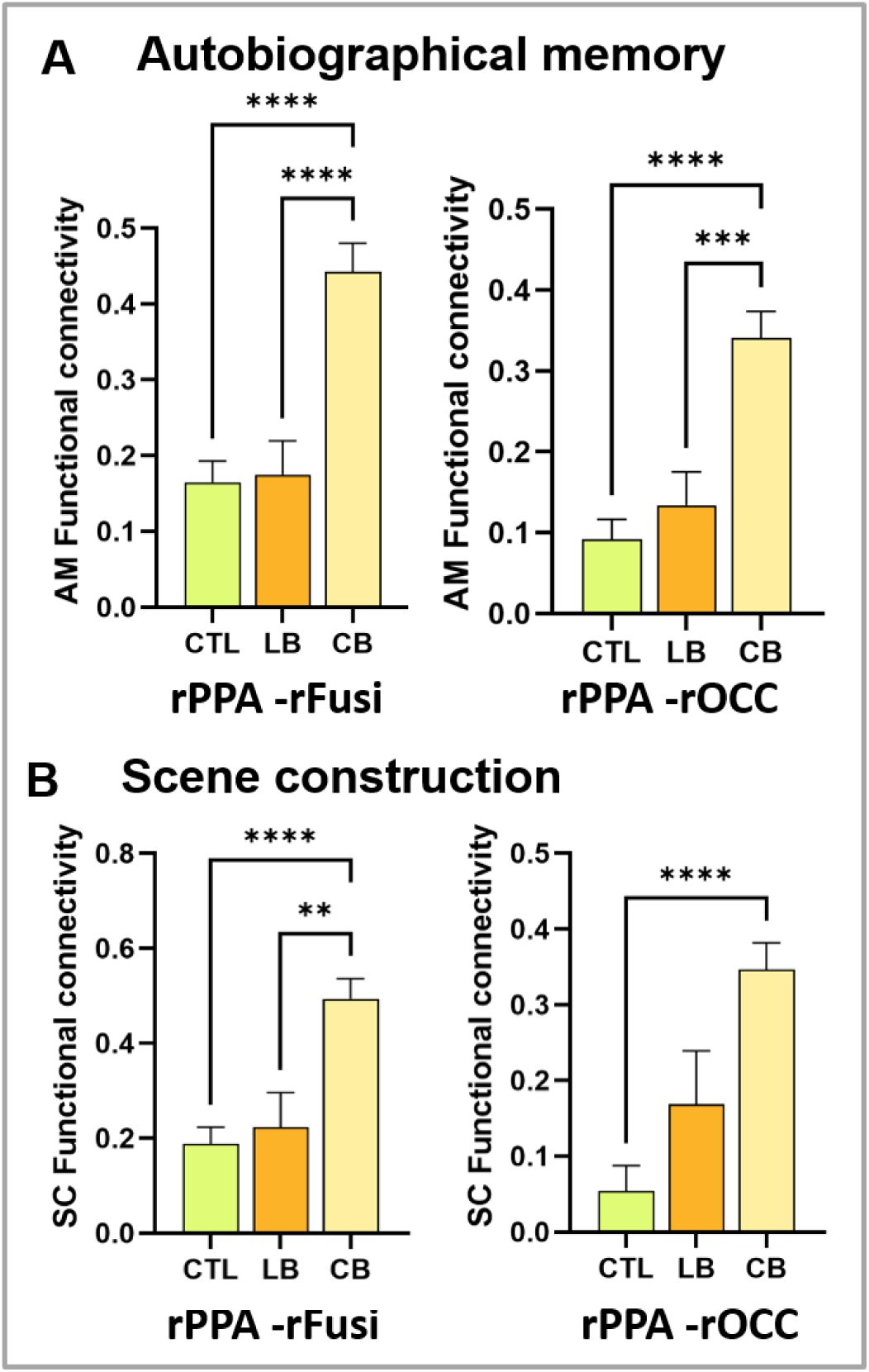
Task-based functional connectivity during autobiographical memory and scene construction. (a) Extracted ROI-to-ROI functional connectivity during autobiographical memory and (b) scene construction for sighted controls (CTL), late-blind (LB), and congenitally blind (CB) participants. CB participants exhibited increased connectivity between the parahippocampal place area (PPA) and occipital cortex, as well as fusiform gyrus, compared to CTL. LB showed intermediate profiles. Bars represent means ± SEM; significance markers indicate ANCOVA results (p < 0.01, *p < 0.001, **p < 0.0001).

### 8.3 Diffusion tensor imaging (DTI)

#### 8.3.1 DTI acquisition

Diffusion tensor imaging (DTI) was acquired with 64 diffusion directions and a b-value of 1000 s/mm². The sequence used a voxel size of 2 mm isotropic, a TR of approximately 9000 ms, and a TE of approximately 89 ms.

#### 8.3.2 DTI preprocessing

Diffusion tensor imaging (DTI) were preprocessed using FSL 6.0.7.15 (https://fsl.fmrib.ox.ac.uk). First, brain extraction was performed with BET^19^. Susceptibility-induced distortions were corrected using TOPUP based on pairs of b0 images acquired with reversed phase-encoding directions. Eddy current artifacts and head motion were then corrected using EDDY, which included outlier replacement and rotation of b-vectors. Diffusion tensors were fitted with DTIFIT to derive voxelwise fractional anisotropy (FA) and mean diffusivity (MD) maps. All diffusion-derived images were visually inspected for artifacts and preprocessing failures prior to analysis.

#### 8.3.3 DTI analysis

Voxelwise analysis was conducted using the Tract-Based Spatial Statistics (TBSS) pipeline. Individual FA images were nonlinearly aligned to the FMRIB58 FA template, resampled to 1×1×1 mm standard space, and projected onto a mean FA skeleton thresholded at FA > 0.2. MD maps were processed using the tbss_non_FA workflow to ensure identical registration and skeleton projection.

Group comparisons were performed using permutation-based general linear models implemented in randomise with 5,000 permutations and TFCE-based family-wise error correction at p < 0.05. Age and sex were included as covariates in all models. In addition to group contrasts (CTL > CB; CTL > LB), separate voxelwise analyses examined associations between FA and Mental Time Travel (MTT) performance within each group. TFCE-corrected clusters were binarized and intersected with the mean FA skeleton, and participant-level mean FA and MD values were extracted for descriptive statistics. MD values were scaled by a factor of 1,000 and reported in units of ×10⁻³ mm²/s.

#### 8.3.4 DTI results

Tract-based spatial statistics revealed widespread reductions in fractional anisotropy (FA) in blind participants compared to sighted controls. For congenitally blind individuals, significant clusters were primarily located in cerebellar–brainstem pathways and extended into long-range association tracts, including the cingulum and inferior fronto-occipital fasciculus. Late-blind participants showed more extensive occipital involvement, with clusters overlapping the optic radiation, vertical occipital fasciculus, and inferior fronto-occipital fasciculus. No significant differences emerged for mean diffusivity (MD) in any contrast.

Within-group analyses indicated that FA correlated positively with Mental Time Travel (MTT) performance in congenitally blind participants. These associations were distributed across commissural and projection pathways, with the largest clusters spanning cingulum subsections and fronto-occipital connections. No significant correlations were observed in late-blind or sighted participants.

**Supplementary Table S3.** White-matter clusters showing significantly increased fractional anisotropy (FA) in healthy participants compared with congenitally blind (CB) individuals, identified using tract-based spatial statistics (TBSS; *p* < 0.05, TFCE-corrected). For each cluster, voxel extent, maximum TFCE-derived probability (MaxP), and peak MNI coordinates are reported. White-matter tract labels and overlap percentages reflect the proportion of each cluster intersecting individual pathways based on the XTRACT HCP probabilistic tract atlas and the JHU ICBM-DTI-81 White-Matter Labels. Cingulum subsections (Temporal, Peri-genual, Dorsal) are listed without overlap values because these subdivisions are not included in the XTRACT HCP or JHU ICBM-DTI-81 atlases queried with *atlasquery*. FA = fractional anisotropy; MTT = middle thalamic tract; TBSS = tract-based spatial statistics; TFCE = threshold-free cluster enhancement; MaxP = maximum TFCE-derived probability; MNI = Montreal Neurological Institute; HCP = Human Connectome Project; JHU ICBM-DTI-81 = Johns Hopkins University International Consortium for Brain Mapping diffusion tensor atlas; L = left; R = right.

| Cluster ID | Cluster size (voxels) | MAXP | MNI Coordinates (X,Y,Z) |  |  | Tract (Overlap %) |
| --- | --- | --- | --- | --- | --- | --- |
| 1 | 2 | 0.95 | 73 | 70 | 43 | Middle Cerebellar Peduncle (14.9) |
| 2 | 23 | 0.95 | 37 | 120 | 92 | Superior Longitudinal Fasciculus R (12.3) |
| 3 | 40 | 0.95 | 69 | 64 | 44 | Corticospinal Tract R (0.4) |
| 4 | 60 | 0.951 | 71 | 77 | 40 | Middle Cerebellar Peduncle (18.8) |
| 5 | 94 | 0.951 | 125 | 122 | 112 | Superior Longitudinal Fasciculus 2 L (21.0) |
| 6 | 122 | 0.952 | 108 | 78 | 39 | Middle Cerebellar Peduncle (13.8) |
| 7 | 367 | 0.953 | 63 | 57 | 39 | Middle Cerebellar Peduncle (7.5) |
| 8 | 681 | 0.954 | 93 | 87 | 32 | Middle Cerebellar Peduncle (12.4), Corticospinal Tract R (2.2) |
| 9 | 34986 | 1 | 55 | 63 | 68 | Cingulum subsection Temporal R, Cingulum subsection Peri-genual R, Cingulum subsection Peri-genual L, Cingulum subsection Dorsal R, Cingulum subsection Dorsal L, Inferior Fronto-Occipital Fasciculus L (5.1), Forceps Major (4.2), Inferior Fronto-Occipital Fasciculus R (4.2), Optic Radiation L (3.6), Optic Radiation R (3.5), Middle Longitudinal Fasciculus R (3.0), Middle Longitudinal Fasciculus L (2.7), Forceps Minor (1.7), Inferior Longitudinal Fasciculus R (1.7), Anterior Thalamic Radiation L (1.7), Uncinate Fasciculus L (1.4), Anterior Thalamic Radiation R (1.4), Superior Longitudinal Fasciculus 3 R (1.2), Vertical Occipital Fasciculus R (1.0), Uncinate Fasciculus R (1.0), Vertical Occipital Fasciculus L (0.9), Acoustic Radiation R (0.9), Arcuate Fasciculus R (0.8), Inferior Longitudinal Fasciculus L (0.8), Superior Thalamic Radiation L (0.7), Corticospinal Tract L (0.7), Frontal Aslant Tract R (0.7), Acoustic Radiation L (0.7), Arcuate Fasciculus L (0.6), Superior Thalamic Radiation R (0.6), Superior Longitudinal Fasciculus 3 L (0.6), Superior Longitudinal Fasciculus 2 L (0.6), Corticospinal Tract R (0.6), Superior Longitudinal Fasciculus 2 R (0.5), Superior Longitudinal Fasciculus 1 R (0.4), Frontal Aslant Tract L (0.2), Fornix L (0.1), Fornix R (0.1), Anterior Commissure (< 0.0), Superior Longitudinal Fasciculus 1 L (< 0.0) |

**Supplementary Table S4.** White-matter clusters showing significantly increased fractional anisotropy (FA) in healthy participants compared with late-blind individuals, identified using tract-based spatial statistics (TBSS; *p* < 0.05, TFCE-corrected). For each cluster, voxel extent, maximum TFCE-derived probability (MaxP), and peak MNI coordinates are reported. White-matter tract labels and overlap percentages reflect the proportion of each cluster intersecting individual pathways based on the XTRACT HCP probabilistic tract atlas and the JHU ICBM-DTI-81 White-Matter Labels. The dorsal cingulum subsection is listed without overlap values because these subdivisions are not included in the XTRACT HCP or JHU ICBM-DTI-81 atlases queried with *atlasquery*. FA = fractional anisotropy; MTT = middle thalamic tract; TBSS = tract-based spatial statistics; TFCE = threshold-free cluster enhancement; MaxP = maximum TFCE-derived probability; MNI = Montreal Neurological Institute; HCP = Human Connectome Project; JHU ICBM-DTI-81 = Johns Hopkins University International Consortium for Brain Mapping diffusion tensor atlas; L = left; R = right.

| Cluster ID | Cluster size (voxels) | MAXP | MNI Coordinates (X,Y,Z) |  |  | Tract (Overlap %) |
| --- | --- | --- | --- | --- | --- | --- |
| 1 | 4 | 0.95 | 57 | 49 | 64 | Vertical Occipital Fasciculus R (26.5), Forceps Major (1.3) |
| 2 | 13 | 0.95 | 58 | 45 | 65 | Vertical Occipital Fasciculus R (48.9), Forceps Major (8.6) |
| 3 | 181 | 0.956 | 57 | 55 | 66 | Vertical Occipital Fasciculus R (18.2), Inferior Fronto-Occipital Fasciculus R (1.3), Forceps Major (0.8) |
| 4 | 3609 | 0.997 | 122 | 63 | 72 | Optic Radiation L (25.9), Inferior Fronto-Occipital Fasciculus L (19.8), Middle Longitudinal Fasciculus L (13.4), Forceps Major (10.2), Acoustic Radiation L (3.7), Vertical Occipital Fasciculus L (3.0), Fornix L (0.4), Inferior Longitudinal Fasciculus L (0.3), Corticospinal Tract L (0.1), Anterior Commissure (0.1), Uncinate Fasciculus L (0.0), Arcuate Fasciculus L (0.0), Superior Thalamic Radiation L (0.0), Superior Longitudinal Fasciculus 1 L (0.0) |
| 5 | 4875 | 0.998 | 58 | 63 | 72 | Cingulum subsection Dorsal R, Optic Radiation R (19.1), Inferior Fronto-Occipital Fasciculus R (16.3), Middle Longitudinal Fasciculus R (11.9), Forceps Major (8.2), Vertical Occipital Fasciculus R (3.6), Acoustic Radiation R (3.4), Inferior Longitudinal Fasciculus R (2.7), Corticospinal Tract R (2.1), Anterior Thalamic Radiation R (1.1), Arcuate Fasciculus R (0.4), Superior Thalamic Radiation R (0.4), Fornix R (0.3), Superior Longitudinal Fasciculus 1 R (< 0.0), Anterior Commissure (< 0.0), Superior Longitudinal Fasciculus 2 R (< 0.0) |

**Supplementary Table S5.**
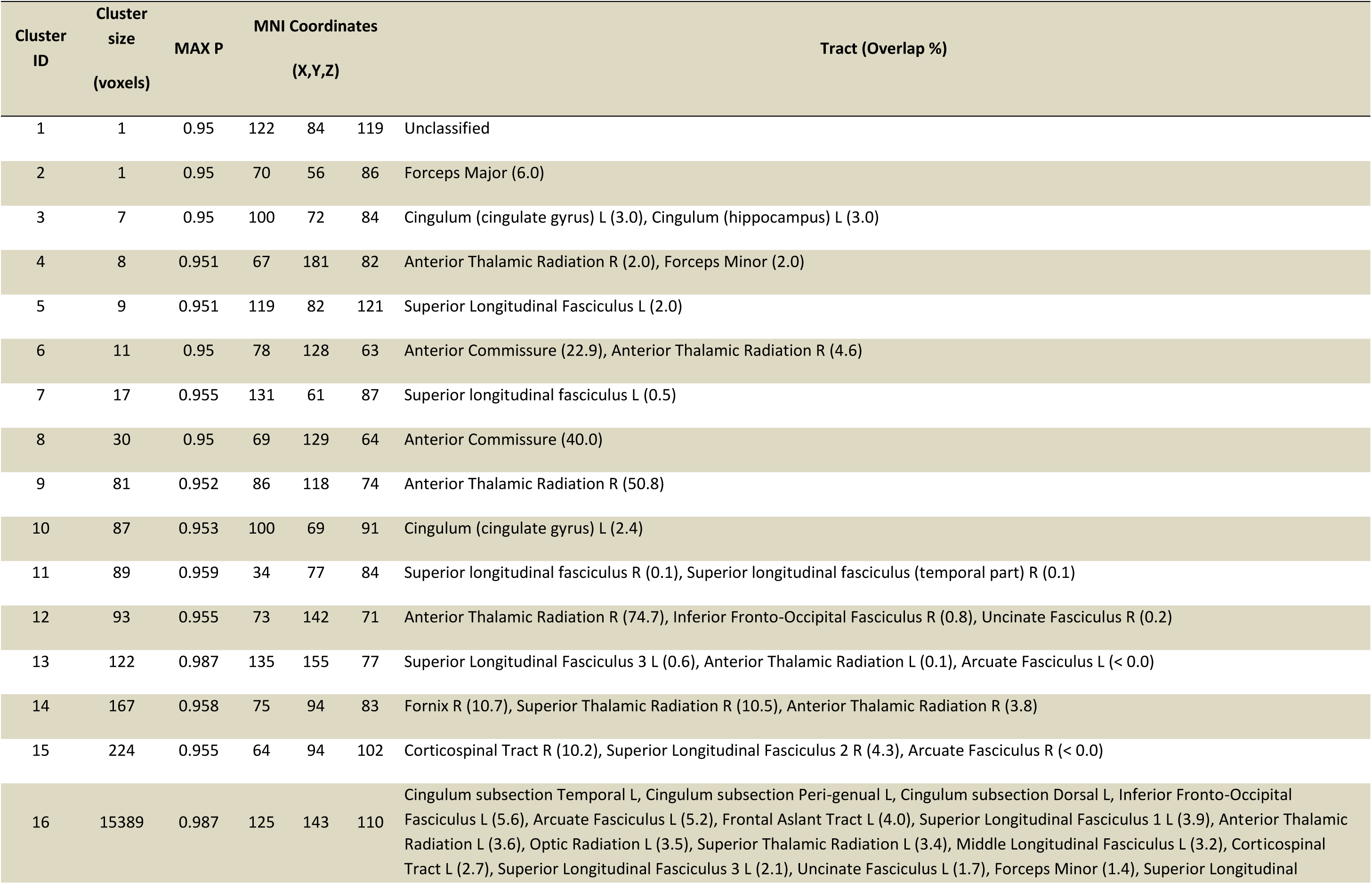
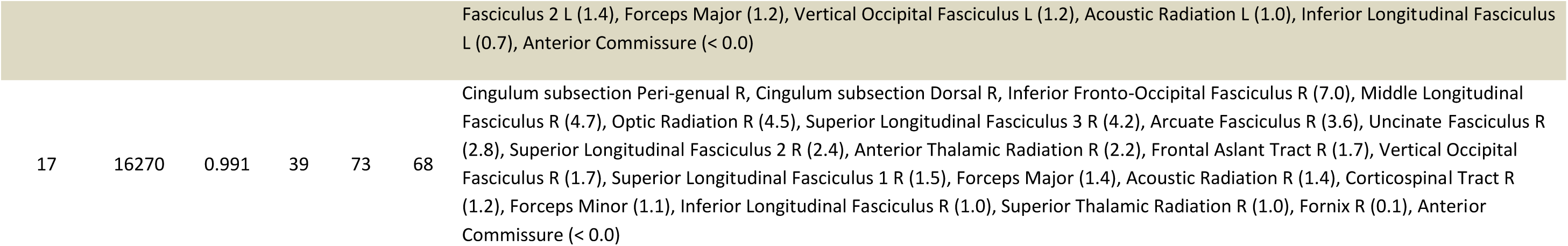
White-matter clusters in congenitally blind (CB) individuals showing significant positive associations between fractional anisotropy (FA) and MTT scores, identified using tract-based spatial statistics (TBSS; p < 0.05, TFCE-corrected). For each cluster, voxel extent, maximum TFCE-derived probability (MaxP), and peak MNI coordinates (x, y, z) are reported. White-matter tract labels and overlap percentages indicate the proportion of cluster voxels intersecting each tract based on the XTRACT HCP probabilistic tract atlas and the JHU ICBM-DTI-81 white-matter labels. Cingulum subsections (temporal, peri-genual, dorsal) are listed without overlap values because these subdivisions are not included in the XTRACT HCP or JHU ICBM-DTI-81 atlases queried with atlasquery. Abbreviations: FA = fractional anisotropy; MTT = middle thalamic tract; TBSS = tract-based spatial statistics; TFCE = threshold-free cluster enhancement; MaxP = maximum TFCE-derived probability; MNI = Montreal Neurological Institute; HCP = Human Connectome Project; JHU ICBM-DTI-81 = Johns Hopkins University International Consortium for Brain Mapping diffusion tensor atlas; L = left; R = right.

### 8.4 Structural volume-based MRI

#### 8.4.1 MRI acquisition

Structural imaging was performed using a multi-echo MPRAGE (MEMPRAGE) sequence with a repetition time (TR) of 2560 ms and four echo times (TEs: 1.72, 3.44, 5.16, and 6.88 ms). Images were acquired at 0.8 mm isotropic resolution with a matrix size of 320 × 320 × 224 and a bandwidth of 680 Hz/pixel. The total acquisition time was approximately 6 minutes and 48 seconds. These MR images were used for whole brain voxel-based morphometry (VBM) analysis and hippocampal segmentation.

#### 8.4.2 Voxel-based morphometry (VBM) processing pipeline

VBM analyses were performed in SPM12 using segmentation into gray matter, white matter, and CSF, followed by high-dimensional registration with the SHOOT template creation algorithm for improved alignment (https://www.fil.ion.ucl.ac.uk/spm/docs/tutorials/vbm/). Normalized gray matter maps were modulated and smoothed with an 6 mm FWHM Gaussian kernel for group-level comparisons. Images were thresholded at a false discovery rate of p<0.05. For display purposes, we then used the ROIs again with the highest fMRI group differences (i.e., rPPA, rFusiform, rOCC, lOCC, and PCC) to extract grey matter volume.

#### 8.4.3 Voxel-based morphometry (VBM) results

VBM revealed widespread group differences in grey matter volume (GMV), primarily within occipital regions and extending into areas associated with scene construction, such as the parahippocampal place area (PPA) and fusiform gyrus (Supplementary Fig. S7). Clusters also overlapped with the medial anterior parahippocampal gyrus typically implicated in spatial and scene-based cognition. These findings suggest that structural adaptations in blindness are concentrated in neocortical regions supporting perceptual and conceptual scaffolds rather than in core episodic hubs.

ROI-based extractions confirmed this pattern. Bar graphs (Supplementary Fig. S4b) illustrate that sighted controls (CTL) exhibited the highest GMV across all regions of interest (PPA, fusiform gyrus, rOCC, lOCC, PCC), followed by congenitally blind participants (CB), with late-blind individuals (LB) showing markedly reduced volumes.

**Supplementary Fig. S8.**
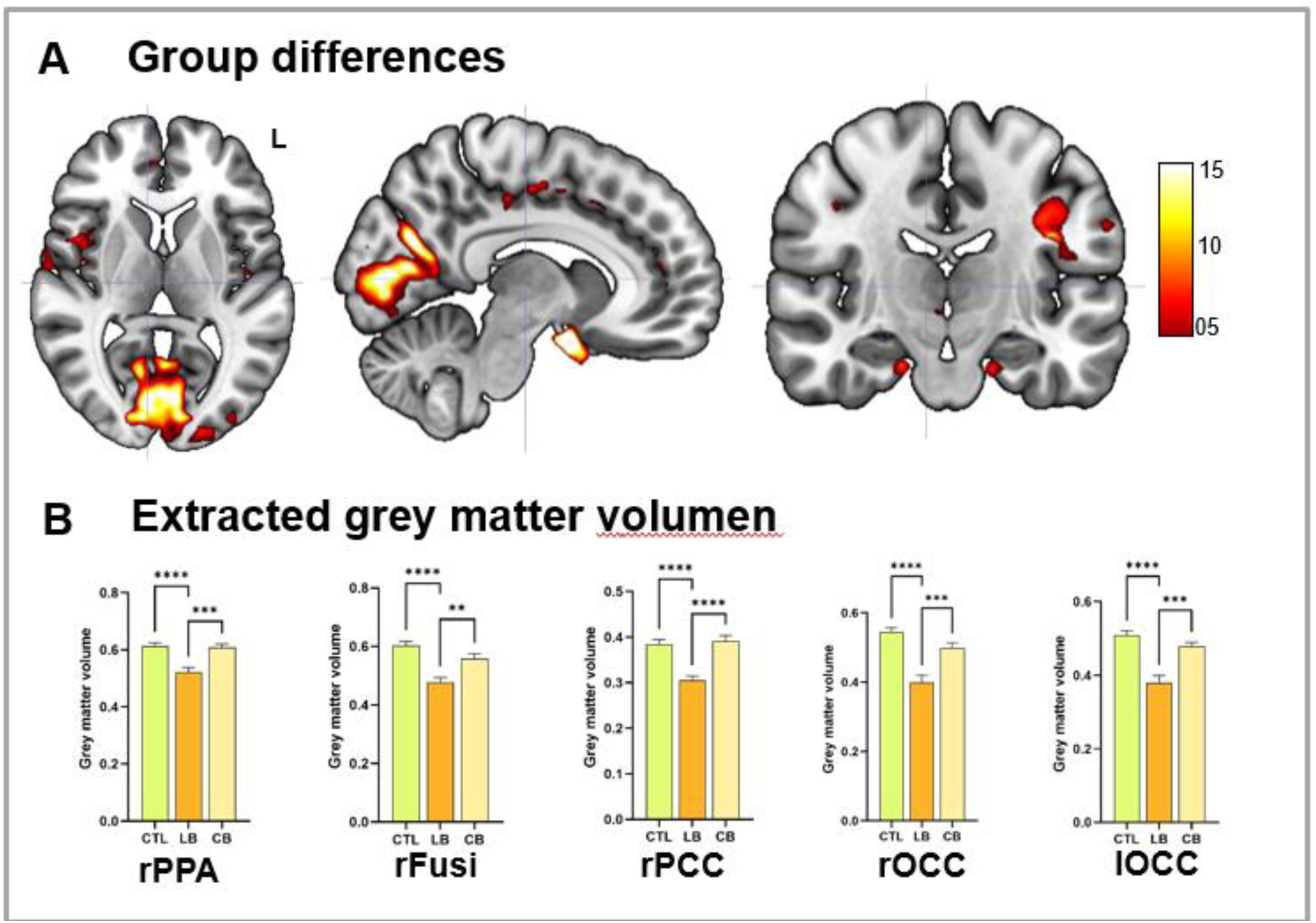
Voxel-based morphometry reveals group differences in gray matter volume. (a) Whole-brain VBM contrasts show significant group differences in gray matter volume (GMV), primarily within occipital regions and extending into areas associated with scene construction, including the parahippocampal place area (PPA). Clusters also overlapped with the medial anterior parahippocampal gyrus, typically implicated in spatial and scene-based cognition. Statistical maps are thresholded at p < 0.05, FDR-corrected, and displayed in MNI152 space; color bar indicates t-values. (b) ROI-based extractions illustrate GMV differences across regions of interest (rPPA, rFusi, rPCC, rOCC, lOCC), confirming that sighted controls (CTL) exhibit the highest GMV, followed by congenitally blind participants (CB), with late-blind individuals (LB) showing markedly reduced volumes.

#### 8.4.4 Hippocampal segmentation processing pipeline

Hippocampal volumes were obtained via FastSurfer automated segmentation with manual correction in ITK-SNAP. Each hippocampus was divided into anterior and posterior portions using the uncus as landmark, and volumes were normalized to total intracranial volume before statistical analysis.Hippocampal volumetry employed FastSurfer for automated segmentation, followed by manual correction in ITK-SNAP. Each hippocampus was divided along its longitudinal axis into anterior and posterior portions using the uncus as anatomical landmark. Volumes were normalized to total intracranial volume prior to statistical analysis. Volumetric analyses were conducted using ordinary one-way ANCOVAs.

#### 8.4.5 Hippocampal segmentation results

An ANCOVA analysis revealed no significant differences in hippocampal volume across groups (F(2, 70) = 1.91, *p* = .16, *n^2^p* = .05). Age had a significant effect on hippocampal volume (F(1, 70) = 5.68, *p* = .02, *n^2^p* = .08). Also, the right and left hippocampi were not statistically different in volume (F2(70) = 2.71, *p* = .073, *n^2^p* =.07; F(2,70) = 1.09, *p* = .34, *n^2^p* = .03). Age had especially a significant impact on right hippocampal volume, F(1, 70)= 9.0, *p* = .004, *n^2^p* = .11.

#### Integrated axis of mental time travel strategies

### 9.1 Principal component analysis

To reduce dimensionality and identify a latent factor summarizing behavioral and neuroimaging measures, we performed a PCA on a curated set of variables from the “PCA” dataset. The included variables were:

- **Behavioural measures:** VVIQ, MTT score (difference between conceptual and perceptual conditions), MTT visual, MTT Non-visusal, and Semantic diversity
- **Neuroimaging measures:** right occipital activation, right parahippocampal activation, left and right hippocampal activation, task-based connectivity, resting-state connectivity, structural connectivity and grey matter volume.

All variables were standardized (z-scored) prior to PCA. We extracted one principal component (PC1) to capture the dominant variance pattern across these measures. PC1 explained 35,1% of the total variance, which is considered substantial for multidimensional behavioral-neuroimaging data.

### 9.2 Component significance

To assess whether PC1 represented a meaningful structure rather than random noise, we conducted a parallel analysis. We generated 1,000 random datasets with identical dimensions and computed their first eigenvalues. The observed eigenvalue for PC1 (3.596) exceeded all random eigenvalues (mean = 1.720), yielding p < 0.0001, confirming that PC1 was statistically significant.

### 9.3 Group comparisons

To test whether PC1 scores differed across experimental groups (Group 1, Group 2, Group 3), we performed a one-way ANOVA. PC1 scores were computed for each participant and sign-adjusted so that Group 3 had positive values and Group 1 negative values for interpretability. Post hoc comparisons were conducted using Tukey’s HSD when the omnibus test was significant.

We found a strong group effect on PC1 scores (F(2, 58) = 80.59, p < 0.0001, η² = 0.779), indicating that group membership explained approximately 73% of the variance in PC1 scores. Tukey post hoc tests confirmed that all pairwise group differences were significant (Group 3 > Group 2 > Group 1, all p < 0.001).

### Statistical analysis

All statistical analyses were performed in SPSS v29.0. Age was included as a covariate in all group comparisons. Behavioral data were analyzed using ANCOVA or MANCOVA for multi-variable outcomes, with Bonferroni-adjusted post hoc tests. Neuroimaging contrasts were assessed using permutation-based GLMs with TFCE correction (p < 0.05), and connectivity analyses applied FDR correction for multiple comparisons.

## Notes

### Competing Interest Statement

The authors have declared no competing interest.

